# Cold exposure drives weight gain and adiposity following chronic suppression of brown adipose tissue

**DOI:** 10.1101/789289

**Authors:** Peter Aldiss, Jo E Lewis, Irene Lupini, Ian Bloor, Ramyar Chavoshinejad, David Boocock, Amanda K Miles, Francis J P Ebling, Helen Budge, Michael E Symonds

## Abstract

Therapeutic activation of thermogenic brown adipose tissue (BAT) may be feasible to prevent, or treat, cardiometabolic disease. However, rodents are commonly housed below thermoneutrality (∼20°C) which can modulate their metabolism and physiology including the hyperactivation of brown (BAT) and beige white adipose tissue. We housed animals at thermoneutrality from weaning to chronically supress BAT, mimic human physiology and explore the efficacy of chronic, mild cold-exposure and β3-adrenoreceptor agonism under these conditions. Using metabolic phenotyping and exploratory proteomics we show that transfer from 28°C to 20°C drives weight gain and a 125% increase in subcutaneous fat mass, an effect not seen with YM-178 administration thus suggesting a direct effect of a cool ambient temperature in promoting weight gain and further adiposity in obese rats. Following chronic suppression of BAT, uncoupling protein 1 mRNA was undetectable in IWAT in all groups. Using exploratory adipose tissue proteomics, we reveal novel gene ontology terms associated with cold-induced weight gain in BAT and IWAT whilst Reactome pathway analysis highlights the regulation of mitotic (i.e. G2/M transition) and metabolism of amino acids and derivatives pathways. Conversely, YM-178 had minimal metabolic-related effects but modified pathways involved in proteolysis (i.e. eukaryotic translation initiation) and RNA surveillance across both tissues. Taken together these findings are indicative of a novel mechanism whereby animals increase body weight and fat mass following chronic suppression of adaptive thermogenesis from weaning. In addition, treatment with a B3-adrenoreceptor agonist did not improve metabolic health in obese animals raised at thermoneutrality.

## Introduction

Therapeutic activation of thermogenic brown adipose tissue (BAT) may be feasible to prevent, or treat, cardiometabolic disease [1]. In rodent models of obesity, the activation of BAT and uncoupling protein (UCP1)-positive beige adipocytes in white adipose tissue (WAT) by cold exposure and sympathomimetics (i.e. β3-agonists) can attenuate or reverse obesity, diabetes and atherosclerosis, thus improving metabolic health [1]. A major factor influencing these outcomes is that animals are typically housed at temperatures well below their thermoneutral zone (which for a rodent is c. 28°C) [2]. Under these conditions, not only is BAT hyperactive, but UCP1+ beige adipocytes are readily seen in the inguinal WAT (IWAT) depot [3].

The ‘cold-stressed’ animal has been widely studied but much less is known about the underlying adaptations in adipose tissue of animals maintained at thermoneutrality. Usually, temperatures as low as 4°C, which represent an ‘extreme cold’, are used to activate BAT, with the induction of UCP1 and subsequent thermogenic response primarily seen in subcutaneous IWAT and other ‘beige’ depots [3]. Conversely, when animals are housed at thermoneutrality and then exposed to 20°C, the induction of UCP1 is primarily seen in BAT. These differences suggest there are two steps to the ‘browning’ process and emphasise the need to study rodent metabolism under thermoneutral conditions. Importantly, it was recently demonstrated that BAT from obese mice, housed chronically at thermoneutrality closely resembles human BAT [4]. This model of ‘physiologically humanised BAT’ is now thought to represent the best choice for studying the physiology of this key metabolic tissue [4]. Here, we extend this recent work by raising animals at thermoneutrality, on an obesogenic diet from weaning. Beginning this early is a particularly important consideration given the early developmental steps which would be occurring in early life [5]. Furthermore, we use rats as classically, their physiology to external stressors such as diet and the environment is closer, than mice, to humans [6]. Using this model we have demonstrated UCP1 mRNA is absent in subcutaneous IWAT (the classical ‘beige’ depot) and not induced with exercise training [7], a common response at standard housing temperatures and one which is typically not seen in humans [8]. We hypothesised that activation of BAT by mild-cold and YM-178, a highly selective β3-agonist that was recently been shown to activate BAT in lean and obese humans, and drive improvements in lipid metabolism, would be negligible under these conditions [9, 10]. Moreover, perivascular BAT would be more responsive as previously shown to be the case with exercise training [7]. Finally, we sought to determine how the AT proteome responds following chronic BAT suppression to better understand the molecular response to potentially thermogenic stimuli at thermoneutrality.

## Results

Four weeks of exposure to 20°C promoted weight gain and increased BAT and subcutaneous IWAT mass, an effect not seen with a clinically relevant dose of YM-178 (Fig. 1A-E). There was no change in total fat mass (i.e. the total of all dissected depots), gonadal (GWAT), mesenteric (MWAT), retroperitoneal (RPWAT) or paracardial (PCAT) fat depots, or in liver or heart mass (Fig. 1C and Supp. Fig 1B-G) suggesting cold does not drive whole body changes in adiposity, and potentially lean mass. There was however a significant increase in kidney size (Supp. Fig 1H). As there was no difference in 24h energy expenditure, or intake, (Supp. Fig. 1I-J) we incorporated these parameters, alongside body mass, into an ANCOVA. Whilst there was no evidence that changes in body mass were associated with altered energy expenditure (Fig. 1G) in cold exposed rats there was a significant, unexpected relationship between body mass and energy intake in this group (r^2^=0.852, p=0.025, Fig. 1H) which was not seen in animals treated with YM-178. (Fig. 1A-H). Increased weight gain and adiposity in cold-exposed animals was not associated with impaired metabolic parameters (i.e. serum glucose, triglycerides and NEFA) or hormones (i.e. insulin and leptin) (Fig. 1I-N) suggesting it is not pathological.

**Figure 1.**
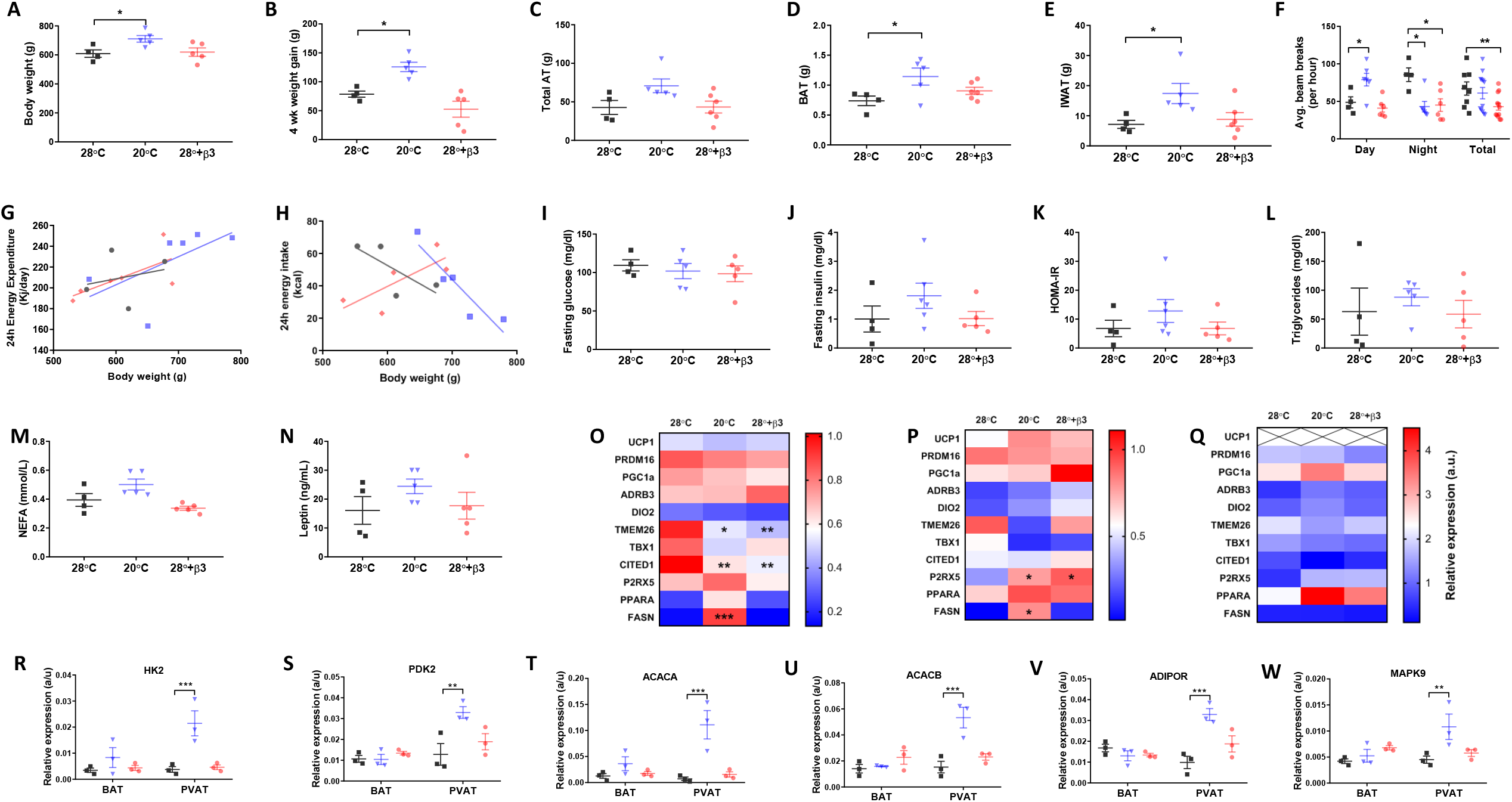
Cold exposure (20°C) but not YM-178 (28°C+β3) drove weight gain and deposition of BAT and inguinal white adipose tissue (IWAT) with no effect on serum metabolites or thermogenic markers. (A) Final body weight, (B) 4 week intervention weight gain, (C) total fat mass, (D) BAT mass, (E) IWAT mass, (F) 24h ambulatory activity, (G) 24h energy expenditure, (H) 24h energy intake, (I-N) serum hormones and metabolites. (O-Q) Markers of brown and beige adipose tissue in BAT, PVAT and IWAT, (R-W) select metabolic genes in BAT and PVAT. Data expressed as mean ± SEM, n=4-5 per group. For comparison, data was analysed by either one (A-E, H-Q), two-way ANOVA (F, R-W) or ANCOVA (G and H) with Sidak post-hoc tests. Significance denoted as * <0.05; ** <0.01 or *** <0.001.

Neither cold exposure, nor YM-178 were effective at inducing thermogenic genes (i.e. UCP1) in BAT or PVAT. in (Fig. 1O-P). Expression of the BAT marker CITED1 and beige marker TMEM26 were reduced in BAT of both cold exposed and YM-178 treated rats whilst the beige marker P2RX5 was upregulated in PVAT. Using targeted arrays to screen for primary genes involved in adipose tissue metabolism we saw an increase in the expression of FASN mRNA in both BAT and PVAT (Fig.1O-P). There was also an increase of genes involved in glycolysis (i.e. HK2 and PDK), fatty acid oxidation (i.e. ACACA and ACACB) and insulin resistance (i.e. AdipoR1 and MAPK9) in PVAT only following cold exposure (Fig. 1R-W). In contrast, in IWAT, UCP1 mRNA (Fig. 1Q).was absent in all rats and, despite a c.125% increase in IWAT mass of cold exposed rats, there was no change in the expression of other genes associated with thermogenesis (i.e. ADRβ3), beige adipocytes (i.e. TMEM26) or lipogenesis (i.e. FASN).

Morphologically, BAT was characterised by a heterogenous mix of small, mitochondria rich lipid droplets and large lipid droplets, and adipocytes, indicative of large-scale whitening (Fig. 2A). Analysis of lipid droplet area in BAT demonstrated a significant increase in both cold, and YM-178 treated rats (Fig. 2B). Adipocyte size also increased in IWAT of cold-exposed rats (Fig. 3A and B). Given the surprising increase in BAT and IWAT mass with cold exposure, we then carried out exploratory adipose tissue proteomics. This method, which quantifies the 30-40% most abundant proteins across samples, did not detect UCP1 in BAT which is not unexpected given the chronic suppression of adaptive thermogenesis. We identified 175 differentially regulated proteins in BAT of cold-exposed rats (Fig. 2C, Table 1 and supp. data) including an increase in the mitochondrial citrate transporter protein (SLC25a1), glucose-6-phosphate dehydrogenase (G6PD), and the muscle isoforms of phosphoglycerate mutase (PGAM2) and creatine kinase (CKm).

**Figure 2.**
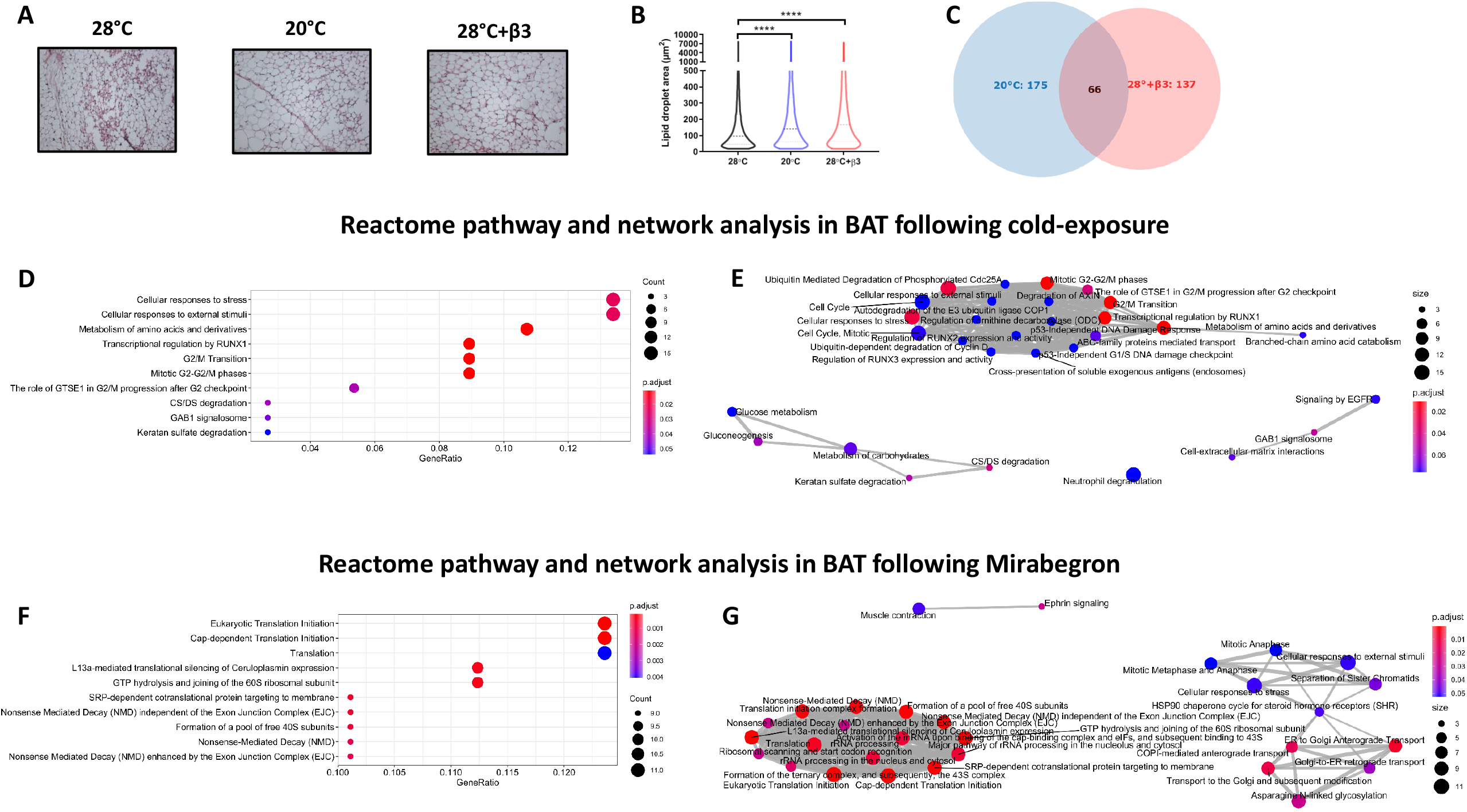
Histological and proteomics analysis of BAT following cold-exposure (20°C) and YM-178 treatment (28°C+β3). (A-B) Histological analysis of BAT and lipid droplet area. (C) Venn diagram of differentially regulated proteins. Reactome pathway analysis detailing enriched pathways and interrelated networks in cold exposed (20°C, D-E) and YM-178 treated animals (28°C+β3, F-G). Data expressed as mean ± SEM, n=4-5 per group. Adipocyte/lipid droplet area quantified using Adiposoft (28°C, n=8949; 20°C, 15512 and 28°C+β3, 12446). For comparison, data was analysed by one-way ANOVA (B) or using the ReactomePA package (D-G). Significance denoted as **** <0.0001.

**Figure 3.**
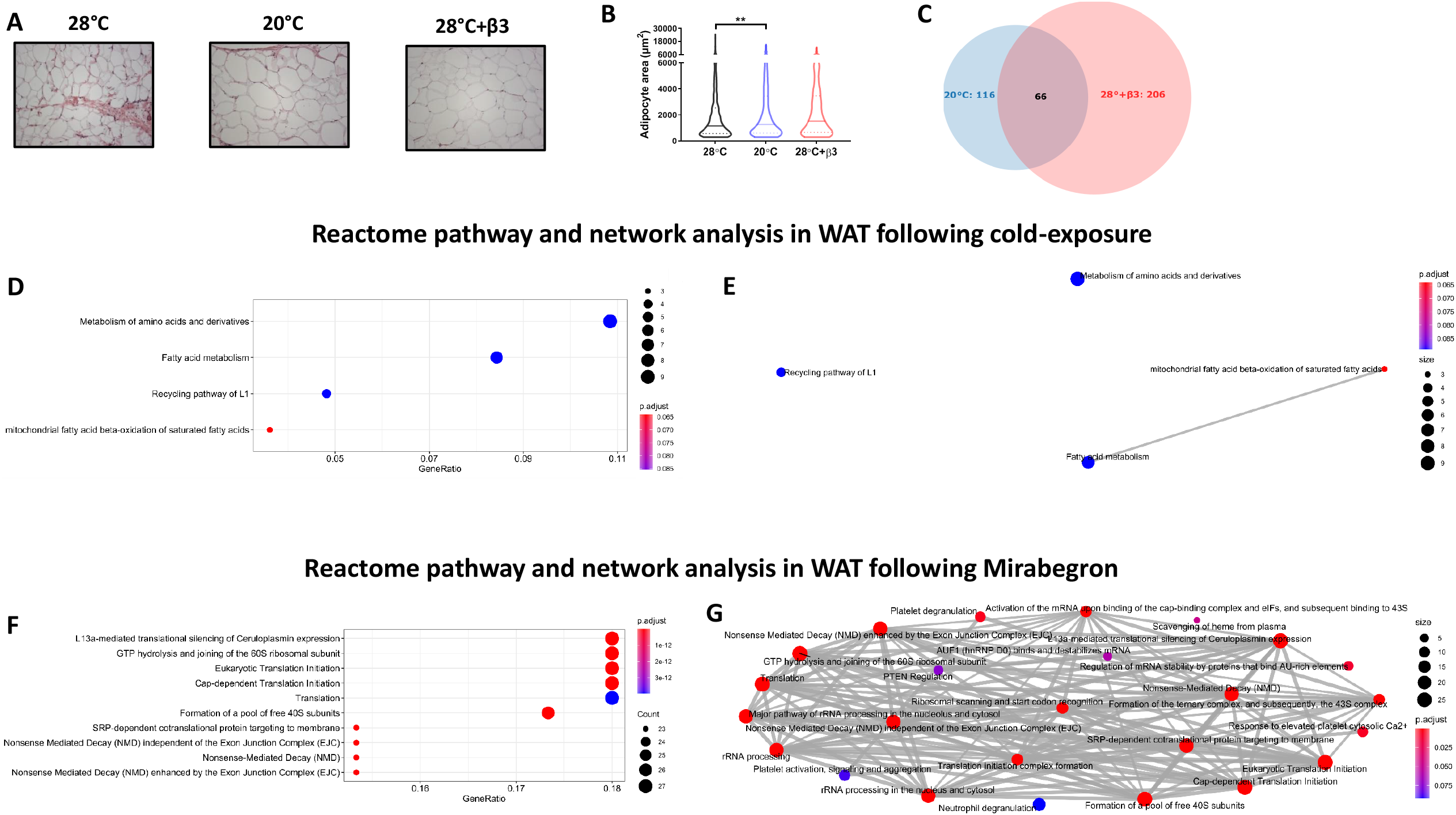
Histological and proteomics analysis of WAT following cold-exposure (20°C) and YM-178 treatment (28°C+β3). (A-B) Histological analysis of BAT and lipid droplet area. (C) Venn diagram of differentially regulated proteins. Reactome pathway analysis detailing enriched pathways and interrelated networks in cold exposed (20°C, D-E) and YM-178 treated animals (28°C+β3, F-G). Data expressed as mean ± SEM, n=4-5 per group. Adipocyte/lipid droplet area quantified using Adiposoft (28°C, n=750; 20°C, 553 and 28°C+β3, 527). For comparison, data was analysed by one-way ANOVA (B) or using the ReactomePA package (D-G). Significance denoted as ** <0.01.

**Table 1.**
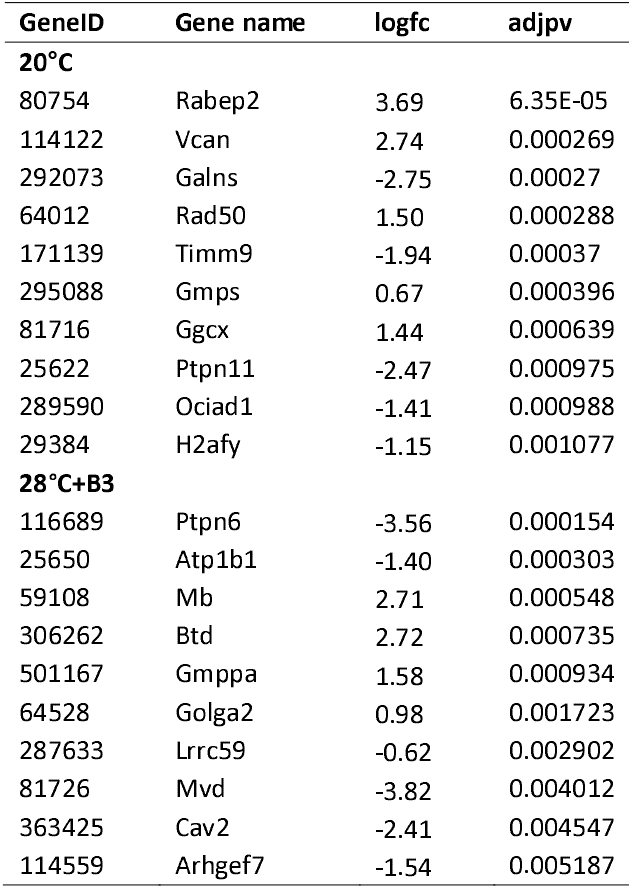
Full list of differentially regulated proteins in BAT

Conversely, 137 proteins were differentially regulated in BAT of YM-178 treated animals (Fig. 2C, Table 1 and supp. data) with an upregulation of proteins involved in skeletal muscle physiology including myosin heavy chain 4 (MYH4), the fast-twitch skeletal muscle isoforms troponin I2 (TNNI2) and calsequestrin 1 (CASQ1in addition to proteins governing endothelial adhesion and vascular growth (PECAM1 and FBLN5). These changes occurred alongside a downregulation of the aldo-keto reductase family member proteins B15 (AKR1B15) and C3 (AKR1C3), mevalonate diphosphate decarboxylase (MVD), trans-2,3-enoyl-CoA reductase (TECR), acyl-CoA dehydrogenase short/branched chain (ACADSB), phosphate cytidylyltransferase 1, choline, alpha (PCYT1A) and 3-hydroxybutyrate dehydrogenase 1 (BDH1) proteins.

We then carried out functional analysis of the BAT proteome. The differentially regulated proteins in BAT of cold exposed animals enriched GO terms involved ‘*glucose import*’, ‘*ATP-dependent helicase activity*’ and ‘*regulation of protein phosphorylation*’ whilst there was also an enrichment of nuclear related GO terms including ‘*histone deacetylation*’, ‘*nucleosomal DNA binding*’, ‘*nuclear chromatin*’ and the ‘*nucleosome*’ (Table 2 and supp. data). Conversely, differentially regulated proteins in BAT of animals treated with YM-178 enriched GO terms including ‘*positive regulation of protein kinase B signalling*’, ‘*negative regulation of cellular carbohydrate metabolic process*’ and ‘*positive regulation of atpase activity*’ whilst there was also an enrichment of GO terms involved in both brown adipocyte and muscle biology including ‘*brown fat cell differentiation*’, ‘*skeletal muscle contraction*’ and ‘*regulation of muscle contraction*’ (Table 2 and supp. data). Finally, using ReactomePA we show enriched pathways regulated by cold exposure (Fig. 2D), and their interactions (Fig. 2E), including ‘transcriptional regulation of RUNx1’, pathways involved in mitosis (i.e. G2/M transition), and the degradation of glycoproteins (i.e. CS/DS degradation). Conversely, YM-178 treatment enriched multiple pathways associated with proteolysis (i.e. eukaryotic translation initiation and formation of a pool of free 40S subunits) and RNA surveillance (i.e. nonsense mediated decay) (Fig. 2F and G).

**Table 2.**
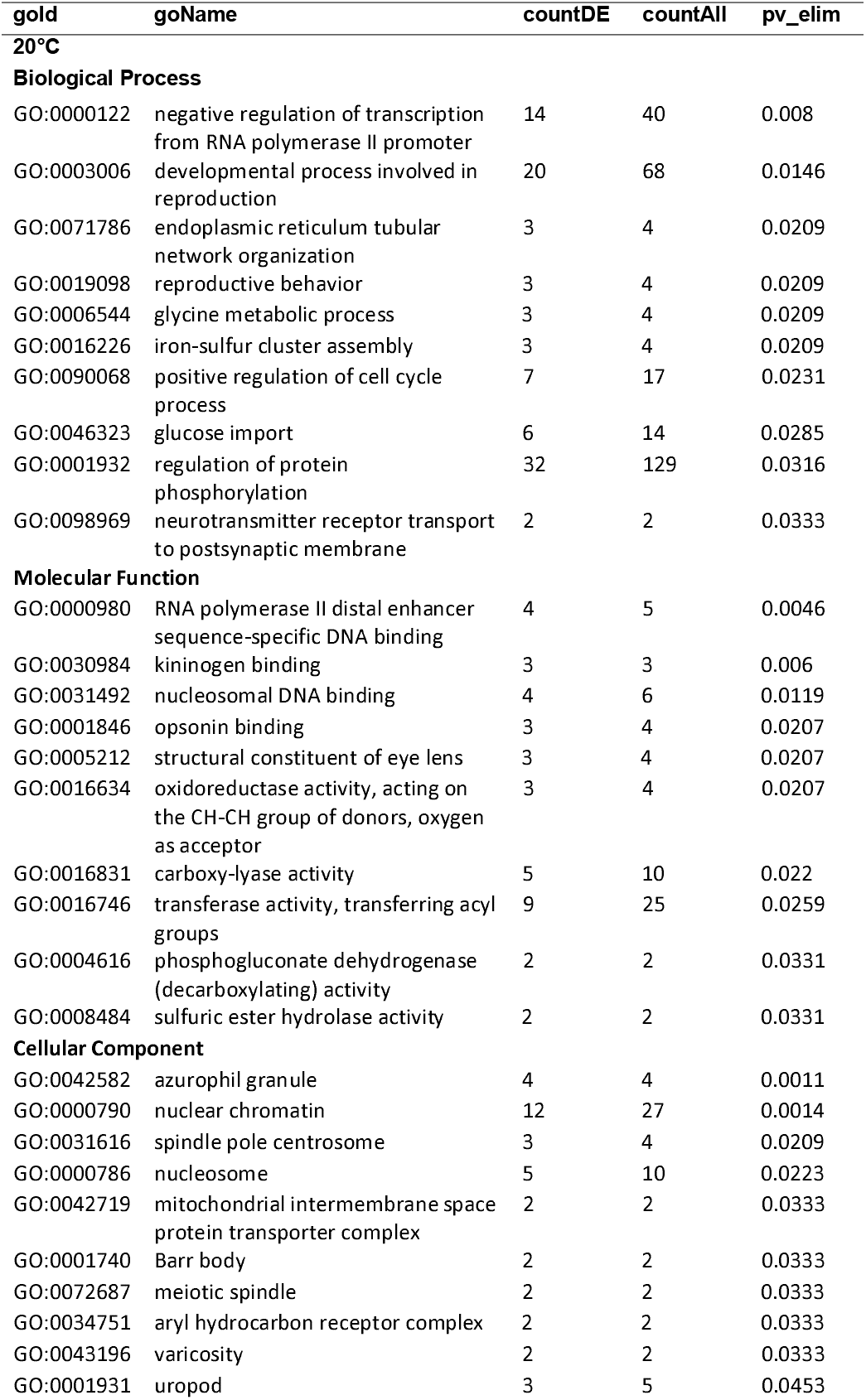
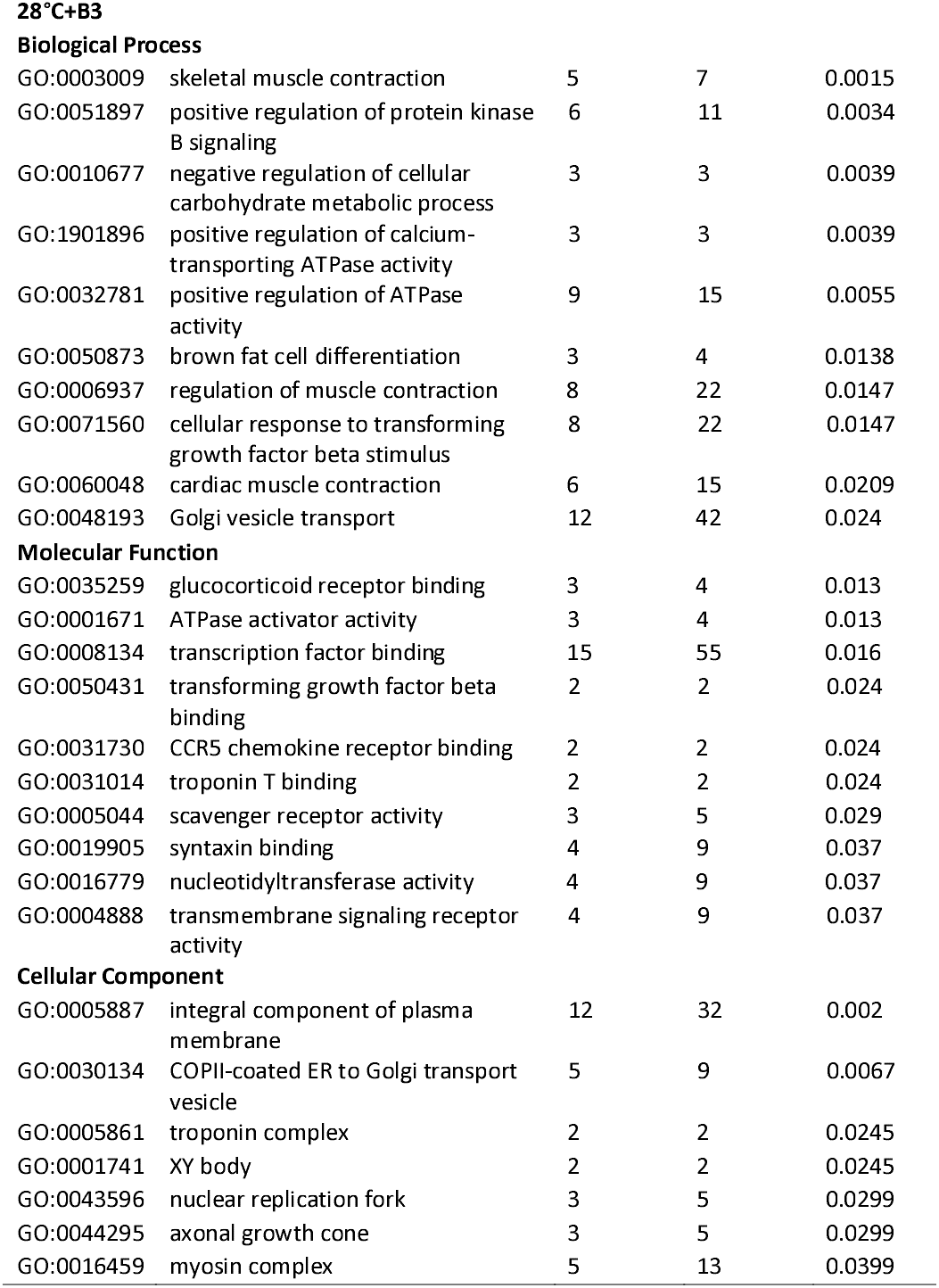
Full list of GO terms enriched in BAT.

The substantial increase in IWAT of cold-exposed rats was associated with 116 differentially regulated proteins (Fig. 3C, Table 3 and supp. data) including the upregulation of lipid metabolic proteins such as fatty acid binding proteins 1 (FABP1) and 3 (FABP3), fatty acid synthase (FASN) and ATP citrate lyase (ACLY) and those involved in beta oxidation including hydroxynacyl-CoA dehydrogenase (HADH) and acyl-CoA dehydrogenase long chain (ACADL). In addition, there was an upregulation of glycerol kinase (GK), pyruvate carboxylase (PC) and monocarboxylic acid transporter 1 (MCT1) that were accompanied by a downregulation of multiple proteins involved in mRNA processing and splicing (i.e. RNA binding motif protein 8B, RBM8B; dexd-Box helicase 39B, DDX39B and poly(u) binding splicing factor 60, PUF60).

**Table 3.**
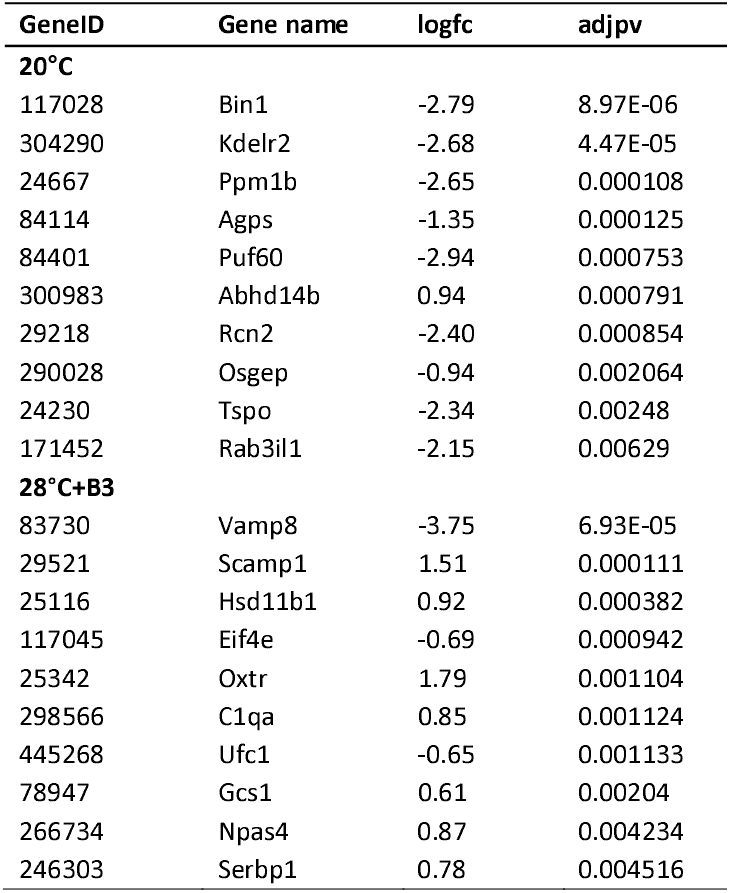
Full list of differentially regulated proteins in WAT

YM-178 administration modulated 206 proteins in IWAT (Fig. 3C, Table 3 and supp. data) including an upregulation of multiple proteins involved in the nervous system including synuclein gamma (SNCG), neurolysin (NLN) and neuronal pas domain protein 4 (NPAS4). This was associated with an increase in proteins involved in lipid and cholesterol metabolism including carnitine palmitoyltransferase 1A (CPT1A), hormone-sensitive lipase (LIPE) and apolipoprotein C1 and M (APOC1 and APOM). Interestingly, YM-178 also induced an increase in multiple inflammatory proteins, including orosomucoid 1 (ORM1), complement C4A (C4A), S100 calcium binding protein A8 (S100A8) and S100 calcium binding protein B (S100B).

Differentially regulated proteins in IWAT of cold-exposed rats enriched GO terms involved in the ‘*DNA damage response’, ‘3-hydroxyacyl-coa dehydrogenase activity*’ and ‘*NAD+ binding*’ whilst YM-178 enriched inflammatory terms including the *‘acute phase response*’ and *‘structural constituent of ribosome*’ suggesting an effect of sympathetic activation on both the inflammatory system and protein synthesis (Table 4 and supp. data). Finally, ReactomePA demonstrated enriched pathways (Fig. 3D), and their interactions (Fig. 3E) were associated with IWAT expansion including the ‘metabolism of amino acids and derivatives’ and ‘fatty acid metabolism’. Conversely, and similar to BAT, YM-178 treatment enriched multiple pathways associated with proteolysis (i.e. eukaryotic translation initiation and formation of a pool of free 40S subunits) and RNA surveillance (i.e. nonsense mediated decay) but to a larger degree (i.e. 23-27 proteins per term rather than 9-11; Fig. 3F and G).

**Table 4.**
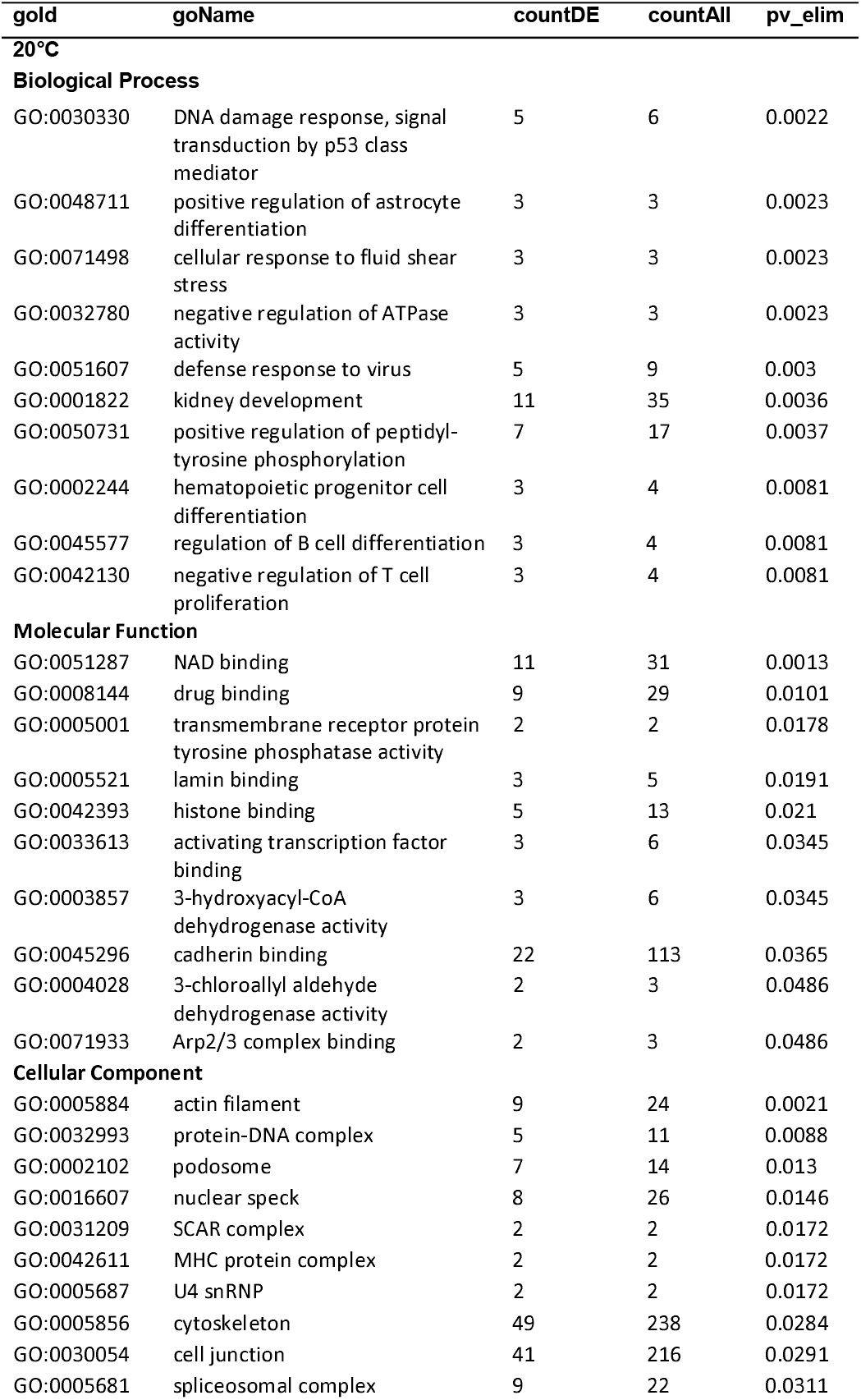
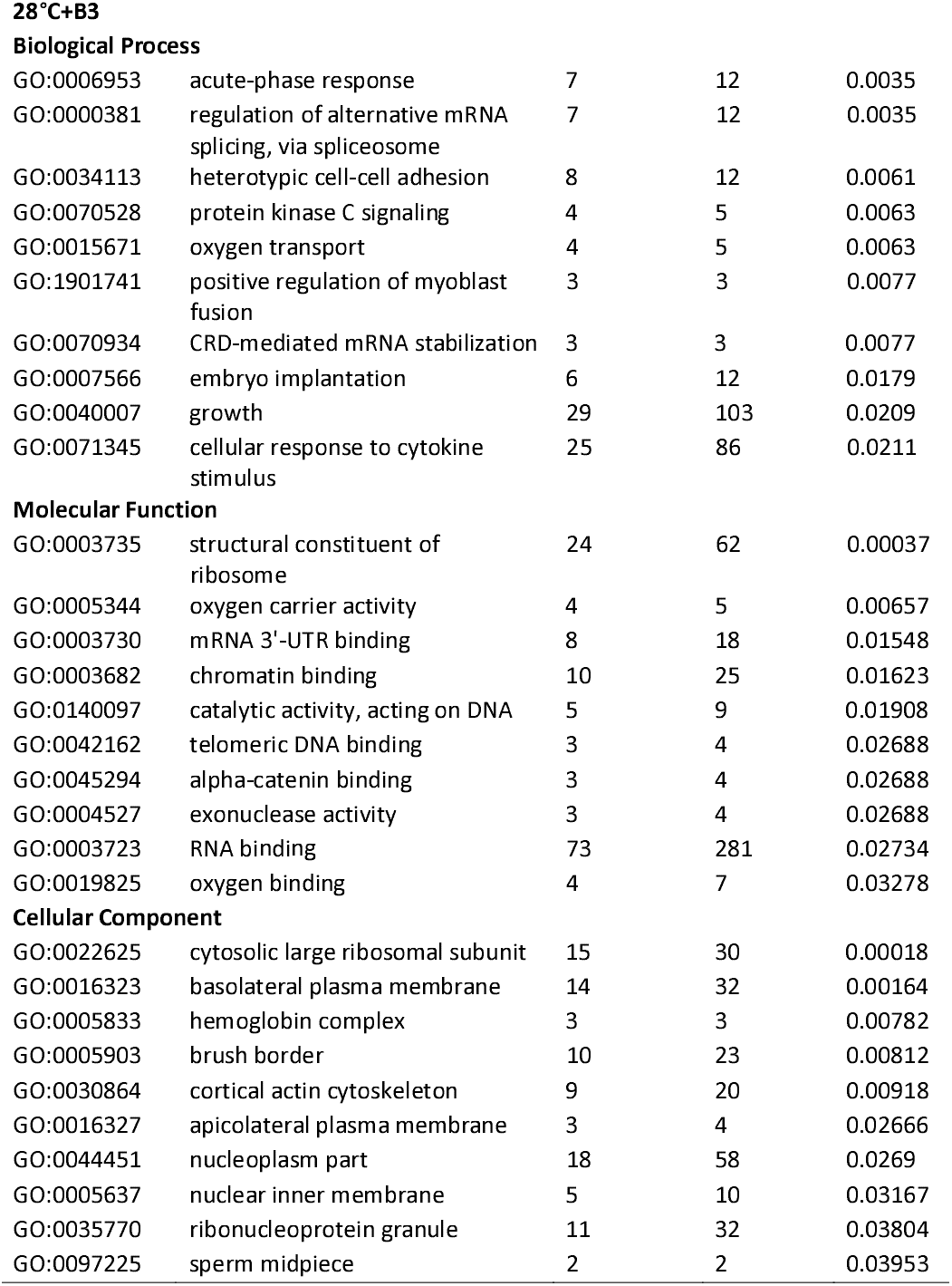
Full list of GO terms enriched in WAT.

## Discussion

Housing temperature, especially cold exposure, impacts on metabolic homeostasis as illustrated by the effects on BAT and ‘browning’ [3, 11-14]. Little is known as to whether interventions considered to promote browning are effective in animals maintained at thermoneutrality [2, 14]. Here we show that chronic exposure to a mild-cold stimulus (i.e. standard housing temperature) drives weight gain and the deposition of large quantities of subcutaneous adipose tissue in obese rats rather than the activation of BAT and subsequent weight-loss as expected [1]. This effect is not seen in rats treated with the highly selective β3-agonist YM-178, suggesting sympathetic activation is not involved and a direct effect of ambient temperature. Taken together these findings are indicative of a novel mechanism whereby rats increase body weight and fat mass following chronic suppression of adaptive thermogenesis from weaning.

There is accumulating evidence that, in the absence of adaptive thermogenesis, other mechanisms compensate in order to maintain body temperature. For instance, in UCP1 k/o mice, a reduction in BAT thermogenesis leads to a the recruitment of shivering thermogenesis in skeletal muscle [15]. Conversely, when shivering is impaired in Sarcolipin k/o mice, there is a compensatory increase in BAT activity [15]. In obese animals lacking BAT, there seems to be an entirely different homeostatic response to cold stress. Following BAT lipectomy, obese, cold-exposed rats gain weight and adipose tissue mass is nearly doubled [16] which mirrors our findings when BAT is chronically supressed. Furthermore, intermittent cold-exposure (from 20°C to 4°C) over a period of days promotes weight gain and adiposity and this is associated with intermittent increases in energy intake [17]. Rats are also more susceptible to weight gain, and fat accumulation when reared in the cold (18°C vs. 30°C), an effect which persists when housed at a common temperature [18]. There is also a plausible mechanism linking the gut to this phenotype which needs to be explored. Cold exposure drives intestinal growth, increased fatty acid absorption and paracellular permeability to nutrients [19, 20]. The efficiency of energy utilisation is also sufficient to maintain core body temperature during acute cold exposure [21]. If the gut can grow, even during periods of energy restriction, and maximise absorption of energy to a degree that it sustains critical functions, and ultimately life, then it may be able to drive adiposity in a similar manner. Whilst the insulative effects of obesity are being debated [22-24] a model whereby rats, and potentially other rodents, deposit subcutaneous fat during exposure to cold (i.e. “store up nuts for the winter internally” [18]) would make sense, and be hugely beneficial from an evolutionary perspective given wild animals cannot simply increase energy intake during winter months.

Another important finding of this study is that the increase in weight gain and subcutaneous AT mass seen in cold-exposed rats was not associated with impaired metabolic parameters (i.e. fasting glucose and lipids) or any discernible adipose tissue dysfunction. Here, chronic exposure to a mild cold stimulus seemingly drives a phenotypically healthy expansion of subcutaneous AT. Using exploratory proteomics, we were able to elucidate processes in both BAT and IWAT associated with this expansion of AT mass. An increase in proteins that modulate metabolism, including those involved in glycolysis, the TCA cycle and lipogenesis, suggests that there is an increased metabolic flux in these depots which, ultimately, results in net lipogenesis. In BAT, this is associated with an enrichment of mitotic pathways (i.e. G2/M transition). The G2/M transition is a critical point in the cell cycle where, following DNA replication, the mitotic process begins, and cells separate into replicate daughter cells. Mitotic clonal expansion is essential for adipogenesis in 3T3-L1 adipocytes [25] and is one of the postulated mechanisms through which FTO regulates fat mass [26]. Further, enrichment of the ‘transcriptional regulation on RUNX1’ pathway, which regulates the white-to-brown/beige transition via CDK6 points towards the cell-cycle as a key mediator of increased BAT mass following cold-exposure [27]. As such, an increase in mitosis of adipocyte precursors, key changes in the cell cycle and subsequent adipogenesis may be partly driving increased BAT mass.

In IWAT, an upregulation of regulatory proteins involved in lipid metabolism (i.e. FABP1, FASN and ACLY) and beta oxidation (i.e. HADH and ACADL) in addition to GK, PC and MCT1 indicates an increased metabolic flux in IWAT. Importantly, however, adipocyte size in IWAT of cold-exposed rats only increased by ∼17%. As such, we predict that this physiological expansion of IWAT is largely due to adipogenesis which ties in to the already established theorem of ‘healthy adipose tissue expansion’ where growth of subcutaneous AT, through adipogenesis, protects from the adverse effects of an obesogenic diet [28].

We were also able to elucidate the impact of sympathetic activation in the absence of any change in housing temperature by administering YM-178. This highly selective β3-agonist increases BAT activity, whole body EE and induces ‘browning’ in humans [29-32]. More recently, two landmark studies have demonstrated that YM-178 also improves glucose homeostasis and insulin sensitivity, improves β-cell function, reduces skeletal muscle triglyceride content and increases HDL cholesterol and adiponectin in overweight and obese prediabetic or insulin resistant humans [9, 10]. However, these, and other studies show that human BAT activity and the response to either cold, or other thermogenic stimuli (i.e. YM-178) is highly heterogenous which is unsurprising given humans undergo seasonal BAT activation to varying degrees [33]. Our aim was to determine the efficacy of this treatment in a homogenous population where BAT had been supressed from early life. Under these conditions, YM-178 treatment had no impact on metabolic parameters or thermogenesis. It does, however, effect pathways involved in protein synthesis in both BAT and, to a larger degree, IWAT. In the hippocampus, β3-AR agonism (by isoprenaline) drives ERK/MTOR dependent activation of eukaryotic initiation factor 4E and inhibition of the translation repressor 4E-BP whilst CL316, 243 and salbutamol drive skeletal muscle hypertrophy and protein synthesis in mice and humans respectively [34-36]. This would suggest a novel role for YM-178 in regulating protein synthesis in adipose tissues which has yet to be examined with previous human studies focussing on overactive bladder and associated symptoms [36, 37]. YM-178 also impacted on proteins involved in the acute phase response and inflammation in both BAT and IWAT. Taken together, this suggests that YM-178 treatment following chronic suppression of BAT may act negatively on AT driving inflammatory processes. Whilst these differences may be species specific, they may also highlight potential effects of YM-178 if given to individuals lacking BAT, or who exhibit low responsiveness to thermogenic stimuli.

Whilst this study points to a novel, unexplained, yet controversial adaptation to cold it is important to remember the major differences in physiology, and cellular processes in animals housed at different ambient temperatures [14]. These differences will likely be exacerbated in animals who have been raised at thermoneutrality from weaning and it is not entirely unexpected that the physiological response to a commonly studied stimulus would be different in this scenario. It is clear that the early life environment, and early life stressors, impact adult physiology and susceptibility to disease, including obesity, and important adaptations during the developmental period may underlie the adaptations seen here. Despite aiming to mimic human physiology this work cannot of course recapitulate the human in-utero, and birth environment and there may be other key developmental changes in humans during this period that are not applicable here. Further, rearing temperature of rats is associated with changes to sympathoadrenal activity and rats are susceptible to obesity in both a strain, and sex-dependent manner [18, 38, 39]. It will be important in future to determine if this effect is not only species specific, but strain and sex specific as we look to untangle aspects of this biology to promote healthy expansion of adipose tissue in humans.

## Conclusion

In summary, we show that chronic exposure to mild-cold following chronic suppression of BAT drives weight gain and the deposition of large quantities of subcutaneous AT. Whilst the precise mechanism is not clear, this work points towards a novel response whereby animals increase body weight and fat mass in response to a reduction in ambient temperature. We propose that chronic suppression of BAT from weaning, using thermoneutral housing and an obesogenic diet, changes the physiological response to cold in the obese state. Instead, humanised animals deposit adipose tissue and gain weight, an effect not seen with YM-178, suggesting a direct effect of temperature, where an insulative mechanism is potentially recruited when a thermogenic response is absent.

## Methods

### Animals, cold exposure and YM-178 treatment

All studies were approved by the University of Nottingham Animal Welfare and Ethical Review Board and were carried out in accordance with the UK Animals (Scientific Procedures) Act of 1986 (PPL no.). Eighteen male Sprague-Dawley rats aged 3 weeks were obtained from Charles River (Kent, UK) and housed immediately at thermoneutrality (c.28°C), on a high-fat diet (HFD; 45%, 824018 SDS, Kent, UK), under a 12:12-hour reverse light-dark cycle (lights off at 08:00). These conditions were chosen so as to closer mimic human physiology [3], minimise animal stress and maximise data quality and translatability [40]. At 12 weeks of age, all animals were randomised to 4 weeks of standard housing temperature (20°C, n=6), YM-178 (28°C+β3, at a clinically relevant dose of 0.75mg/kg/day, n=6) administration or HFD controls (28°C, n=6). In adherence to the National Centre for the Replacement, Refinement and Reduction of Animals in Research (NC3Rs), this experiment was ran alongside our work looking at the effect of exercise training on ‘browning’ and utilised the same cohort of control HFD animals [41].

### Metabolic assessment and tissue collection

All animals were placed in an open-circuit calorimeter (CLAMS: Columbus Instruments, Linton Instrumentation, UK) for the final 48h to enable the assessment of whole body metabolism [42]. All animals were then weighed and fasted overnight prior to euthanasia by rising CO_2_ gradient. BAT, perivascular BAT (PVAT) from the thoracic aorta and IWAT were then rapidly dissected, weighed, snap-frozen in liquid nitrogen and stored at −80°C for subsequent analysis.

### Histology

Adipose tissue histology was performed as previously described [7]. Briefly, BAT and IWAT were fixed in formalin, embedded in wax using an Excelsior ES processor (Thermo-Fisher) and stained using haematoxylin and eosin (Sigma-Aldrich). Between 3 and 5 random sections were then imaged at 10x with an Olympus BX40 microscope and adipocyte size was quantified using Adiposoft [43].

### Gene expression analysis

Total RNA was extracted from each fat depot using the RNeasy Plus Micro extraction kit (Qiagen, West Sussex, UK) using an adapted version of the single step acidified phenol-chloroform method. RT-qPCR was carried out as previously described using rat-specific oligonucleotide primers (Sigma) or FAM-MGB Taqman probes [42]. Gene expression was determined using the GeNorm algorithm against two selected reference genes; *RPL19:RPL13a in BAT* and IWAT (stability value M = 0.163 in BAT and 0.383 in IWAT) and RPL19:HPRT1 in PVAT (stability value M = 0.285).

### Serum analysis

Glucose (GAGO-20, Sigma Aldrich, Gillingham, UK), triglycerides (LabAssay Trigylceride, Wako, Neuss, Germany), non-esterified fatty acids (NEFA-HR(2), Wako, Neuss, Germany), insulin (80-INSRT-E01, Alpco, Salem, NH, USA) and leptin (EZRL-83K, Merck, Darmstadt, Germany) were measured as previously described following manufacturer’s instructions [7].

### Adipose tissue proteomics

Protein extraction, clean up and trypsinisation was performed on 50-100 mg of frozen tissue (n=4/group) was homogenised in 500 μL CellLytic MT cell lysis buffer (Sigma, C3228) prior to removal of lipid and other contaminants using the ReadyPrep 2D cleanup Kit (Biorad, 1632130) [42]. Samples were then subjected to reduction, alkylation and overnight trypsinisation following which they were dried down at 60°C for 4 h and stored at 80°C before resuspension in LCMS grade 5% acetonitrile in 0.1% formic acid for subsequent analysis. Analysis by mass spectrometry was carried out on a SCIEX TripleTOF 6600 instrument [44] with samples analysed in both SWATH (Data Independent Acquisition) and IDA (Information Dependent Acquisition) modes for quantitation and spectral library generation respectively. IDA data was searched together using ProteinPilot 5.0.2 to generate a spectral library and SWATH data was analysed using Sciex OneOmics software [45] extracted against the locally generated library as described previously [42].

### Statistical analysis

Statistical analysis was performed in GraphPad Prism version 8.0 (GraphPad Software, San Diego, CA). Data are expressed as Mean±SEM with details of specific statistical tests in figure legends. Despite prior use of the control group to understand the impact of exercise training only the groups included in this paper were utilised for analyses [7]. Functional analysis of the proteome was performed using the Advaita Bioinformatic iPathwayGuide software (www.advaitabio.com/ipathwayguide.html) (fold change ± 0.5 and confidence score cut-off of 0.75). Significantly impacted biological processes and molecular functions were analysed in the context of the Gene Ontology Consortium database (2017-Nov) [46]. The Elim pruning method, which removes genes mapped to a significant GO term from more general (higher level) GO terms, was used to overcome the limitation of errors introduced by considering genes multiple times [47]. Pathway analysis was carried out using the ReactomePA package on R Studio (version 3.6.2) with a false discovery rate of <0.1.

## Availability of data and material

The datasets used and analysed during the current study are available from the corresponding author on reasonable request.

## Authors’ contributions

P.A., H.B. and M.E.S. conceived the study and attained the funding; P.A. and M.E.S. developed and designed the experiments; P.A, J.E.L, I.L, A.K.M, I.B, R.C and D. J. B. performed the experiments; P.A., A.K.M. and D.J.B. analyzed the data; P.A. and M.E.S. wrote the paper which was revised critically by D.J.B., H.B., F.J.P.E, and J.E.L. for important intellectual content. All authors read and approved the final manuscript.

## Figure legends

**Supplementary Figure 1.**
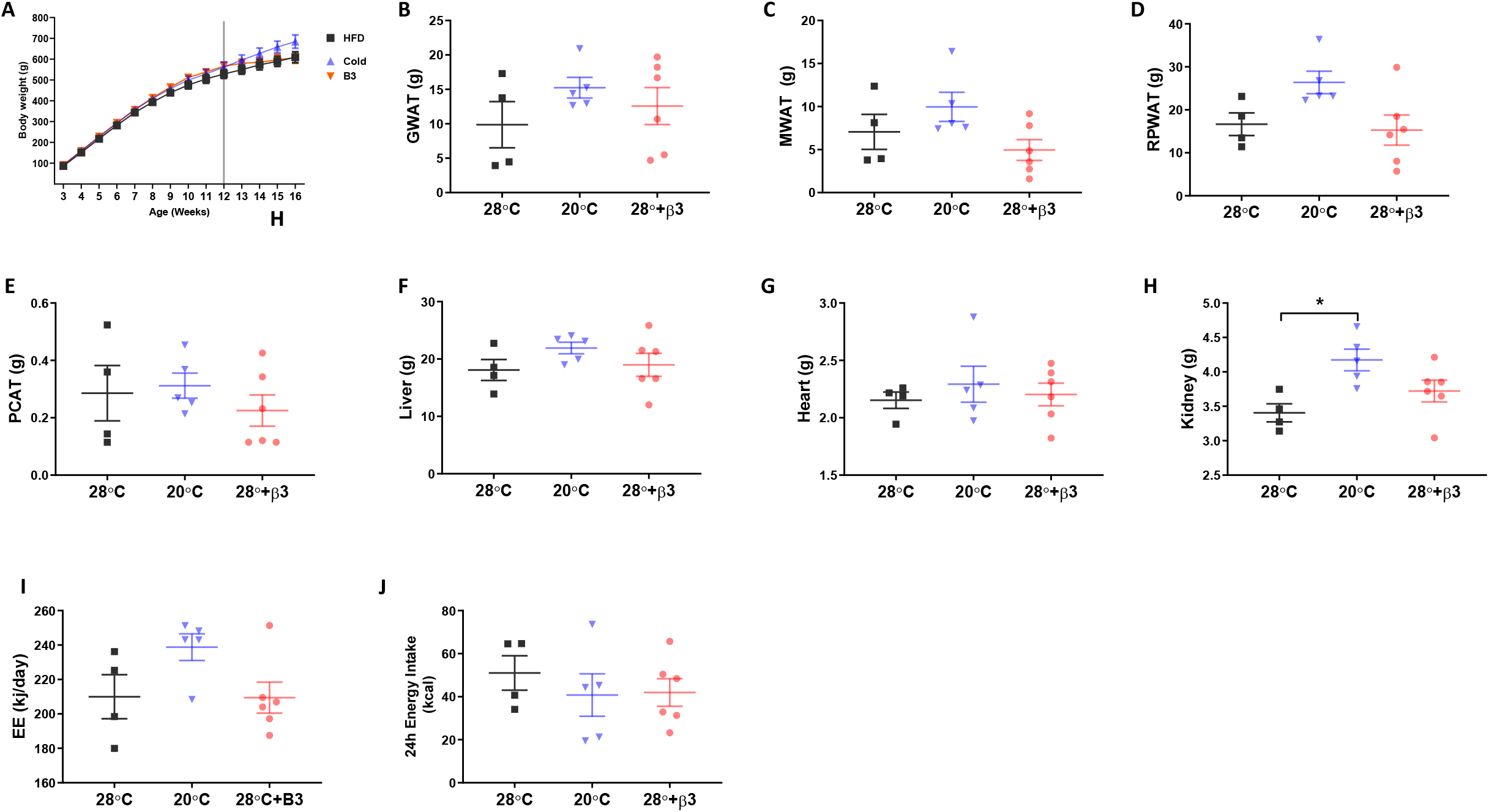
Weight gain trajectory, adipose tissue depot and organ weights. (A) Body weight gain trajectory from weaning and (B) gonadal, (C) mesenteric, (D) retroperitoneal, (E) paracardial adipose tissues. (F) liver, (G) heart and (H) kidney mass. (I-J) absolute energy expenditure and energy intake.

