## Supplementary data for "Cold exposure drives weight gain and adiposity following chronic suppression of brown adipose tissue"

1 *The Early Life Research Unit, Division of Child Health, Obstetrics and Gynaecology, School of Medicine, University of Nottingham;**;**;**;**;*
2 *Nottingham Digestive Disease Centre and Biomedical Research Unit, School of Medicine, University of Nottingham*
3 *School of Life Sciences, Queen’s Medical Centre, University of Nottingham;**;*
4 *School of Biosciences and Veterinary Medicine, University of Camerino, Camerino, MC, Italy;*
5 *John van Geest Cancer Research Centre, Nottingham Trent University, Nottingham, NG11 8N;**;*

| **Table 1.** Full list of differentially regulated proteins in BAT | | | |
| --- | --- | --- | --- |
| **GeneID** | **Gene name** | **logfc** | **adjpv** |
| **20°C** |  |  |  |
| 80754 | Rabep2 | 3.69 | 6.35E-05 |
| 114122 | Vcan | 2.74 | 0.000269 |
| 292073 | Galns | -2.75 | 0.00027 |
| 64012 | Rad50 | 1.50 | 0.000288 |
| 171139 | Timm9 | -1.94 | 0.00037 |
| 295088 | Gmps | 0.67 | 0.000396 |
| 81716 | Ggcx | 1.44 | 0.000639 |
| 25622 | Ptpn11 | -2.47 | 0.000975 |
| 289590 | Ociad1 | -1.41 | 0.000988 |
| 29384 | H2afy | -1.15 | 0.001077 |
| 302669 | Ca5b | 1.18 | 0.001237 |
| 499991 | Steap4 | -3.88 | 0.00228 |
| 315265 | Twf1 | -1.54 | 0.00248 |
| 25283 | Gclc | 2.57 | 0.002571 |
| 84474 | Ddx1 | 2.23 | 0.00326 |
| 94342 | Bag6 | -1.26 | 0.0036 |
| 89827 | Ddx39a | 1.26 | 0.00397 |
| 64679 | Tgm4 | 0.95 | 0.00616 |
| 64517 | Thop1 | 1.25 | 0.006186 |
| 29743 | Slc25a1 | 1.10 | 0.006388 |
| 64630 | Snap23 | -0.57 | 0.006562 |
| 79449 | Rpl21 | -2.43 | 0.007106 |
| 57341 | Parva | -1.36 | 0.009375 |
| 305178 | Hnrnpdl | -0.75 | 0.00987 |
| 85274 | Prdx4 | -3.77 | 0.010514 |
| 259275 | Ostf1 | 0.84 | 0.010579 |
| 29688 | Minpp1 | -2.07 | 0.013155 |
| 445268 | Ufc1 | -2.04 | 0.014068 |
| 24959 | Pgam2 | 1.80 | 0.014105 |
| 361613 | Ppme1 | 2.03 | 0.014231 |
| 307039 | Rab18 | -0.80 | 0.015098 |
| 304024 | Cpox | 2.64 | 0.017902 |
| 100909840 | LOC100909840 | -0.60 | 0.017981 |
| 81781 | Snrpn | -0.51 | 0.018985 |
| 690131 | Hist2h2aa2 | -0.61 | 0.019641 |
| 497198 | Impact | -1.80 | 0.020479 |
| 307905 | Usp10 | -3.13 | 0.020618 |
| 499839 | RGD1564664 | -2.70 | 0.021669 |
| 25010 | Scgb2a1 | -3.44 | 0.022101 |
| 308650 | Hnrnph2 | 2.88 | 0.022881 |
| 310695 | Kirrel1 | 2.62 | 0.025104 |
| 363013 | Tmem123 | 2.59 | 0.026566 |
| 170751 | Xpnpep1 | 3.29 | 0.02881 |
| 100360180 | Pgd | 0.53 | 0.028867 |
| 25420 | Cryab | 0.90 | 0.029983 |
| 311422 | Itpa | 0.78 | 0.032824 |
| 29676 | Psmb3 | -1.65 | 0.033466 |
| 29734 | Hspa13 | 0.88 | 0.034403 |
| 25537 | Rock2 | 2.20 | 0.036381 |
| 305497 | Cobl | 2.18 | 0.03795 |
| 290640 | Map1s | -3.42 | 0.042638 |
| 29637 | Hmgcs1 | 4.22 | 0.042914 |
| 308384 | Sae1 | -2.63 | 0.043105 |
| 300757 | Hexa | 1.15 | 0.046193 |
| 83764 | Flot2 | 1.95 | 0.048583 |
| 103689947 | LOC103689947 | -0.50 | 0.055615 |
| 25425 | Ctsh | 0.58 | 0.055862 |
| 140868 | Fabp5 | 0.63 | 0.057439 |
| 50681 | Acox1 | -1.28 | 0.058114 |
| 295692 | Nup35 | 0.61 | 0.063838 |
| 314648 | Ncln | -4.13 | 0.063902 |
| 64157 | Ddah1 | 1.36 | 0.066755 |
| 81666 | Gnaq | -1.16 | 0.068826 |
| 192360 | Eml2 | 0.81 | 0.07351 |
| 303518 | Smarce1 | 4.10 | 0.073866 |
| 307503 | Etf1 | -1.94 | 0.074281 |
| 192235 | Hyou1 | 1.79 | 0.074523 |
| 287828 | Jpt1 | 0.79 | 0.074615 |
| 297566 | Atp6v1e1 | 0.62 | 0.074767 |
| 84357 | Sh3kbp1 | 1.45 | 0.075562 |
| 117104 | Ppp2r2a | 0.71 | 0.075654 |
| 192249 | Ehd3 | 1.65 | 0.075682 |
| 58815 | Glrx3 | -3.87 | 0.077933 |
| 117028 | Bin1 | -0.89 | 0.082843 |
| 59303 | Tmem33 | 1.06 | 0.085507 |
| 29425 | Psmb5 | 2.58 | 0.08587 |
| 24157 | Acaa1a | -0.72 | 0.089194 |
| 306283 | Anxa8 | -0.66 | 0.089575 |
| 362015 | Ampd2 | 1.19 | 0.090078 |
| 691657 | Crip1 | 1.32 | 0.094208 |
| 294673 | Hexb | -2.92 | 0.102142 |
| 79210 | Fstl1 | 1.71 | 0.102491 |
| 680522 | Hist1h1b | 1.38 | 0.103409 |
| 683788 | Fscn1 | 0.53 | 0.106285 |
| 64032 | Ctgf | -0.50 | 0.106913 |
| 100361558 | LOC100361558 | 1.24 | 0.107837 |
| 114766 | Phb2 | -1.11 | 0.108469 |
| 29254 | Mgll | 0.74 | 0.112464 |
| 287042 | Nubp1 | 1.49 | 0.114342 |
| 64203 | Bcat2 | -1.08 | 0.114377 |
| 102550391 | LOC102550391 | -0.71 | 0.114543 |
| 361532 | Sirt2 | 1.80 | 0.115482 |
| 361635 | LOC361635 | -1.12 | 0.120085 |
| 157074 | Sdha | 0.55 | 0.122822 |
| 64347 | Sncg | 0.94 | 0.132378 |
| 361884 | Mccc2 | -0.58 | 0.13291 |
| 298609 | Efhd2 | 1.97 | 0.132936 |
| 89841 | Pcyt2 | 2.41 | 0.133672 |
| 25277 | Mfge8 | 1.81 | 0.133882 |
| 27139 | Rps26 | -0.91 | 0.134129 |
| 83781 | Lgals3 | 0.70 | 0.13543 |
| 299027 | Eif2s3 | 0.53 | 0.136792 |
| 361092 | Stk24 | 1.58 | 0.138388 |
| 360820 | Pxn | 2.40 | 0.138502 |
| 29153 | Capn1 | 1.18 | 0.140233 |
| 287125 | Nubp2 | -1.48 | 0.142311 |
| 89825 | Nap1l1 | -2.20 | 0.144302 |
| 85492 | Psmb7 | 2.27 | 0.147263 |
| 63938 | Hibadh | -0.71 | 0.152486 |
| 60373 | Nop58 | 2.02 | 0.153349 |
| 64538 | Ilkap | -1.24 | 0.153505 |
| 301252 | Hsp90ab1 | -0.69 | 0.154211 |
| 25177 | Fhl1 | 1.57 | 0.154475 |
| 252928 | Timm13 | -1.25 | 0.154815 |
| 54321 | Cnn3 | 2.50 | 0.155197 |
| 362634 | C1qc | -1.45 | 0.155978 |
| 300981 | Acy1 | -0.60 | 0.15657 |
| 117041 | Nln | -2.71 | 0.161382 |
| 24655 | Plcd1 | 1.36 | 0.16585 |
| 25344 | Phb | -0.91 | 0.166831 |
| 300968 | Uba5 | 2.00 | 0.167161 |
| 25380 | Anxa1 | 0.91 | 0.170097 |
| 315707 | Csk | -3.52 | 0.170238 |
| 282827 | Aip | 0.99 | 0.171773 |
| 114123 | Sardh | 0.82 | 0.172193 |
| 114113 | Pafah1b3 | -3.72 | 0.173067 |
| 108348260 | LOC108348260 | -1.77 | 0.17431 |
| 301618 | Ppp1r7 | 2.07 | 0.176101 |
| 81661 | Gmfb | -3.15 | 0.179404 |
| 361927 | Fxr1 | 1.22 | 0.182166 |
| 315664 | Kdelc2 | -3.19 | 0.182306 |
| 297893 | Hdac1 | 1.93 | 0.185012 |
| 117130 | Grifin | -1.17 | 0.187328 |
| 60466 | Stx7 | 1.26 | 0.191458 |
| 313770 | Mxra8 | 1.83 | 0.191508 |
| 29318 | Ddt | -0.85 | 0.191745 |
| 291081 | Tubb2b | 0.53 | 0.195288 |
| 314644 | Dohh | -2.00 | 0.196706 |
| 24437 | H1f0 | 5.60 | 0.197351 |
| 24787 | Sod2 | -0.79 | 0.197772 |
| 29332 | Stmn1 | -3.43 | 0.198118 |
| 60384 | Copb2 | -1.96 | 0.198454 |
| 501232 | Cesl1 | 1.17 | 0.198671 |
| 29681 | C1qbp | -0.75 | 0.199398 |
| 362282 | Pck1 | -0.86 | 0.199468 |
| 298370 | Txndc12 | -0.69 | 0.199832 |
| 298441 | Nasp | -1.22 | 0.200112 |
| 24377 | G6pd | 0.71 | 0.201778 |
| 317259 | Nono | -0.71 | 0.202927 |
| 58835 | Phgdh | 0.60 | 0.203119 |
| 297699 | Strap | 3.00 | 0.204399 |
| 83572 | Pafah1b1 | -0.64 | 0.204626 |
| 25581 | Psmc2 | -2.54 | 0.205136 |
| 64201 | Slc25a11 | -4.80 | 0.207437 |
| 25643 | Gnai3 | 2.86 | 0.207957 |
| 286896 | Sgpl1 | 1.24 | 0.209271 |
| 246298 | Retsat | 0.60 | 0.209863 |
| 691947 | Eif3j | 1.23 | 0.211981 |
| 171155 | Hadhb | -0.80 | 0.212642 |
| 64665 | Flot1 | 1.11 | 0.216866 |
| 79248 | Abca2 | 0.82 | 0.218028 |
| 293719 | Ubxn1 | 1.55 | 0.220695 |
| 65204 | Cnn1 | 5.54 | 0.223415 |
| 24265 | Ckm | 2.78 | 0.223581 |
| 287876 | Actg1 | 0.54 | 0.229322 |
| 117103 | Rab8a | 2.01 | 0.229323 |
| 299201 | Dlst | -0.60 | 0.230346 |
| 140934 | Elp1 | 0.69 | 0.23234 |
| 313047 | Yars | 1.50 | 0.233809 |
| 171133 | Gcsh | -1.65 | 0.23408 |
| 140931 | Hnrnph1 | -1.11 | 0.235857 |
| 290401 | Esd | 1.41 | 0.238483 |
| 66028 | Arl6ip5 | 3.95 | 0.238743 |
| 310635 | Arhgef2 | -2.38 | 0.239455 |
| 25697 | Ctsl | 1.23 | 0.23982 |
| **28°C+B3** |  |  |  |
| 116689 | Ptpn6 | -3.56 | 0.000154 |
| 25650 | Atp1b1 | -1.40 | 0.000303 |
| 59108 | Mb | 2.71 | 0.000548 |
| 306262 | Btd | 2.72 | 0.000735 |
| 501167 | Gmppa | 1.58 | 0.000934 |
| 64528 | Golga2 | 0.98 | 0.001723 |
| 287633 | Lrrc59 | -0.62 | 0.002902 |
| 81726 | Mvd | -3.82 | 0.004012 |
| 363425 | Cav2 | -2.41 | 0.004547 |
| 114559 | Arhgef7 | -1.54 | 0.005187 |
| 25737 | Pcna | 1.41 | 0.005462 |
| 171516 | Akr1c3 | -3.61 | 0.006199 |
| 311328 | Rmdn3 | -2.54 | 0.006373 |
| 312398 | Smarcad1 | -2.97 | 0.006491 |
| 29583 | Pecam1 | 3.43 | 0.006613 |
| 25491 | Nes | -1.35 | 0.008957 |
| 25106 | Rgn | 1.79 | 0.009092 |
| 117099 | Bdh1 | -1.76 | 0.009659 |
| 100364457 | LOC100364457 | 0.98 | 0.010813 |
| 681429 | Rps27l | 1.68 | 0.011212 |
| 24439 | Hagh | 4.52 | 0.01207 |
| 365377 | Trim72 | -2.27 | 0.012326 |
| 266605 | Dcps | 0.74 | 0.012836 |
| 113922 | Selenof | -1.84 | 0.013927 |
| 81922 | Sh3gl1 | -0.70 | 0.015702 |
| 116463 | Akr1b7 | -4.12 | 0.016705 |
| 309593 | Gnl1 | 1.21 | 0.017056 |
| 360543 | Myh4 | 3.19 | 0.021934 |
| 294853 | Krt18 | -4.53 | 0.022348 |
| 291434 | Rpl17 | 0.57 | 0.024257 |
| 266759 | Hspa4 | 1.93 | 0.025477 |
| 29671 | Psma4 | 2.04 | 0.027228 |
| 100910732 | LOC100910732 | -4.65 | 0.02735 |
| 29437 | Acta1 | 2.41 | 0.031493 |
| 295703 | Serping1 | 1.24 | 0.033653 |
| 25618 | Acadsb | -2.31 | 0.034211 |
| 81651 | Cspg4 | 1.02 | 0.034735 |
| 124461 | Pacsin2 | -1.11 | 0.036395 |
| 260321 | Fkbp4 | -1.82 | 0.039467 |
| 364064 | Pycr2 | 1.40 | 0.040704 |
| 681913 | Gstz1 | 1.68 | 0.041137 |
| 170845 | Ndel1 | -5.37 | 0.041759 |
| 116547 | S100a8 | -1.40 | 0.045185 |
| 366734 | Bag5 | -3.35 | 0.04572 |
| 296570 | Edf1 | -2.35 | 0.046418 |
| 81520 | Marcksl1 | -0.88 | 0.046841 |
| 140544 | Pcyt1a | -2.97 | 0.047611 |
| 501203 | Myl12a | 0.52 | 0.049269 |
| 83712 | Rbbp7 | -2.70 | 0.051168 |
| 25371 | Adprh | -1.34 | 0.052742 |
| 498545 | Tsc22d1 | -2.36 | 0.056873 |
| 108348287 | LOC108348287 | 1.58 | 0.05858 |
| 29435 | Ssr4 | 2.47 | 0.061261 |
| 25484 | Myo1e | -2.31 | 0.062683 |
| 287191 | Rars | 3.19 | 0.067376 |
| 81815 | Tpp2 | -2.81 | 0.069101 |
| 84379 | Rab6a | 0.51 | 0.070121 |
| 59114 | Slc9a3r1 | 1.19 | 0.078356 |
| 140673 | Napa | 1.34 | 0.07892 |
| 313647 | Hp1bp3 | 6.26 | 0.079022 |
| 298098 | Pole3 | 0.74 | 0.079152 |
| 362154 | Zc3h15 | -1.20 | 0.08213 |
| 25338 | Ninj1 | -2.78 | 0.082898 |
| 25621 | Cd81 | -0.60 | 0.084793 |
| 81761 | Rnpep | 1.43 | 0.086835 |
| 361207 | Tmed9 | 2.00 | 0.089368 |
| 114861 | Scpep1 | 0.83 | 0.089463 |
| 65028 | Dnaja1 | -1.01 | 0.089869 |
| 59107 | Ltbp1 | -2.60 | 0.092309 |
| 292023 | Aars | -0.73 | 0.092548 |
| 29285 | Rps15 | -2.97 | 0.096805 |
| 29283 | Rpl29 | 0.70 | 0.097067 |
| 24946 | F9 | 1.73 | 0.098183 |
| 191576 | Tecr | -2.74 | 0.098231 |
| 296320 | Ctnnbl1 | -1.42 | 0.10886 |
| 289456 | Hsd17b11 | 0.92 | 0.110043 |
| 686019 | Casq1 | 2.72 | 0.110524 |
| 100359982 | Mpc2 | -5.03 | 0.113167 |
| 29158 | Fbln5 | 2.97 | 0.114676 |
| 314730 | Ikbip | -1.76 | 0.116982 |
| 246325 | Kcnh8 | 0.84 | 0.117573 |
| 58945 | Dynll1 | 5.47 | 0.118062 |
| 408248 | Psma3l | -0.90 | 0.119604 |
| 361730 | Tkfc | -2.20 | 0.12089 |
| 259246 | LOC259246 | -0.81 | 0.124788 |
| 360629 | Nt5c3b | -1.17 | 0.126636 |
| 684527 | Crtc1 | 2.24 | 0.127565 |
| 171063 | Gtf3c1 | -1.91 | 0.130115 |
| 94266 | Rps27 | 0.77 | 0.136434 |
| 288022 | Ccdc50 | -2.18 | 0.143163 |
| 192269 | Sub1 | 2.53 | 0.143954 |
| 362173 | Caprin1 | -3.02 | 0.144588 |
| 299923 | Ndrg1 | -1.10 | 0.148883 |
| 114023 | Copb1 | 1.04 | 0.155732 |
| 116482 | Sacm1l | 2.79 | 0.155855 |
| 25291 | Anxa3 | -0.51 | 0.159588 |
| 117268 | Khdrbs1 | -3.10 | 0.165319 |
| 619580 | Ctps2 | -0.62 | 0.166527 |
| 94174 | Tinagl1 | -3.84 | 0.166646 |
| 298566 | C1qa | 1.55 | 0.171369 |
| 25475 | Lgals5 | 2.84 | 0.171861 |
| 313200 | Hsdl2 | 0.77 | 0.17566 |
| 25073 | Scarb1 | 1.21 | 0.176283 |
| 296851 | Pon2 | -0.64 | 0.179464 |
| 113956 | Pecr | 2.01 | 0.181245 |
| 308796 | Mesd | -1.28 | 0.183137 |
| 289144 | Cacybp | -0.55 | 0.183169 |
| 171105 | Lnpep | -1.78 | 0.183999 |
| 94197 | Rab14 | 2.32 | 0.188069 |
| 64367 | Ppib | -0.58 | 0.191775 |
| 25125 | Stat3 | -2.20 | 0.19204 |
| 245955 | Lgals3bp | -1.76 | 0.194232 |
| 301442 | Sumo1 | -0.98 | 0.194283 |
| 171562 | Ero1a | 0.75 | 0.196251 |
| 81827 | Psmc5 | -2.94 | 0.199308 |
| 25614 | Ptk2 | 1.49 | 0.201321 |
| 290500 | Ggact | 2.29 | 0.204519 |
| 93646 | Sec31a | 1.75 | 0.211889 |
| 54318 | Eif2s1 | -2.50 | 0.216204 |
| 296554 | Tubb4b | -0.53 | 0.216562 |
| 266760 | Nalcn | -3.14 | 0.217229 |
| 81763 | Rpl5 | 2.72 | 0.222544 |
| 84401 | Puf60 | -2.17 | 0.224326 |
| 29389 | Tnni2 | 2.61 | 0.224664 |
| 117152 | Cand1 | 0.74 | 0.224995 |
| 140922 | Txnl1 | -0.68 | 0.231544 |
| 24614 | Orm1 | 0.85 | 0.233416 |
| 117272 | Prpsap2 | 1.91 | 0.233601 |
| 338401 | Crip2 | 1.21 | 0.237137 |
| 64196 | Safb | 0.53 | 0.238084 |
| 116666 | Lman1 | -0.57 | 0.242846 |
| 64667 | Sgta | -0.88 | 0.242935 |
| 81776 | Rps24 | 1.18 | 0.24314 |
| 84471 | Snx1 | -2.20 | 0.243648 |
| 64198 | Pmpcb | -1.07 | 0.246348 |
| 171145 | Eif2b3 | 2.20 | 0.248176 |
| 81653 | Dbn1 | -1.67 | 0.248852 |

| **Table 2. Full list of GO terms enriched in BAT** | | | | |
| --- | --- | --- | --- | --- |
| **goId** | **goName** | **countDE** | **countAll** | **pv_elim** |
| **20°C** | | | | |
| **Biological Process** | | | | |
| GO:0000122 | negative regulation of transcription from RNA polymerase II promoter | 14 | 40 | 0.008 |
| GO:0003006 | developmental process involved in reproduction | 20 | 68 | 0.0146 |
| GO:0071786 | endoplasmic reticulum tubular network organization | 3 | 4 | 0.0209 |
| GO:0019098 | reproductive behavior | 3 | 4 | 0.0209 |
| GO:0006544 | glycine metabolic process | 3 | 4 | 0.0209 |
| GO:0016226 | iron-sulfur cluster assembly | 3 | 4 | 0.0209 |
| GO:0090068 | positive regulation of cell cycle process | 7 | 17 | 0.0231 |
| GO:0046323 | glucose import | 6 | 14 | 0.0285 |
| GO:0001932 | regulation of protein phosphorylation | 32 | 129 | 0.0316 |
| GO:0098969 | neurotransmitter receptor transport to postsynaptic membrane | 2 | 2 | 0.0333 |
| GO:1903044 | protein localization to membrane raft | 2 | 2 | 0.0333 |
| GO:0032962 | positive regulation of inositol trisphosphate biosynthetic process | 2 | 2 | 0.0333 |
| GO:0032986 | protein-DNA complex disassembly | 2 | 2 | 0.0333 |
| GO:0050884 | neuromuscular process controlling posture | 2 | 2 | 0.0333 |
| GO:0051482 | positive regulation of cytosolic calcium ion concentration involved in phospholipase C-activating G-protein coupled signaling pathway | 2 | 2 | 0.0333 |
| GO:1902667 | regulation of axon guidance | 2 | 2 | 0.0333 |
| GO:0031498 | chromatin disassembly | 2 | 2 | 0.0333 |
| GO:0019322 | pentose biosynthetic process | 2 | 2 | 0.0333 |
| GO:0060766 | negative regulation of androgen receptor signaling pathway | 2 | 2 | 0.0333 |
| GO:0060338 | regulation of type I interferon-mediated signaling pathway | 2 | 2 | 0.0333 |
| GO:1990592 | protein K69-linked ufmylation | 2 | 2 | 0.0333 |
| GO:0003085 | negative regulation of systemic arterial blood pressure | 2 | 2 | 0.0333 |
| GO:2000757 | negative regulation of peptidyl-lysine acetylation | 2 | 2 | 0.0333 |
| GO:1900138 | negative regulation of phospholipase A2 activity | 2 | 2 | 0.0333 |
| GO:0045039 | protein import into mitochondrial inner membrane | 2 | 2 | 0.0333 |
| GO:0006689 | ganglioside catabolic process | 2 | 2 | 0.0333 |
| GO:2000322 | regulation of glucocorticoid receptor signaling pathway | 2 | 2 | 0.0333 |
| GO:0071169 | establishment of protein localization to chromatin | 2 | 2 | 0.0333 |
| GO:0061582 | intestinal epithelial cell migration | 2 | 2 | 0.0333 |
| GO:0009051 | pentose-phosphate shunt, oxidative branch | 2 | 2 | 0.0333 |
| GO:0070193 | synaptonemal complex organization | 2 | 2 | 0.0333 |
| GO:0070192 | chromosome organization involved in meiotic cell cycle | 2 | 2 | 0.0333 |
| GO:0008277 | regulation of G-protein coupled receptor protein signaling pathway | 5 | 11 | 0.0349 |
| GO:0006334 | nucleosome assembly | 6 | 15 | 0.0405 |
| GO:0046578 | regulation of Ras protein signal transduction | 6 | 15 | 0.0405 |
| GO:0010501 | RNA secondary structure unwinding | 4 | 8 | 0.0413 |
| GO:0007626 | locomotory behavior | 7 | 19 | 0.043 |
| GO:0043901 | negative regulation of multi-organism process | 7 | 19 | 0.043 |
| GO:0072321 | chaperone-mediated protein transport | 3 | 5 | 0.0452 |
| GO:0071468 | cellular response to acidic pH | 3 | 5 | 0.0452 |
| GO:0051785 | positive regulation of nuclear division | 3 | 5 | 0.0452 |
| GO:0045599 | negative regulation of fat cell differentiation | 3 | 5 | 0.0452 |
| GO:0031647 | regulation of protein stability | 15 | 53 | 0.0455 |
| GO:1901653 | cellular response to peptide | 16 | 58 | 0.0492 |
| **Molecular Function** | |  |  |  |
| GO:0000980 | RNA polymerase II distal enhancer sequence-specific DNA binding | 4 | 5 | 0.0046 |
| GO:0030984 | kininogen binding | 3 | 3 | 0.006 |
| GO:0031492 | nucleosomal DNA binding | 4 | 6 | 0.0119 |
| GO:0001846 | opsonin binding | 3 | 4 | 0.0207 |
| GO:0005212 | structural constituent of eye lens | 3 | 4 | 0.0207 |
| GO:0016634 | oxidoreductase activity, acting on the CH-CH group of donors, oxygen as acceptor | 3 | 4 | 0.0207 |
| GO:0016831 | carboxy-lyase activity | 5 | 10 | 0.022 |
| GO:0016746 | transferase activity, transferring acyl groups | 9 | 25 | 0.0259 |
| GO:0004616 | phosphogluconate dehydrogenase (decarboxylating) activity | 2 | 2 | 0.0331 |
| GO:0008484 | sulfuric ester hydrolase activity | 2 | 2 | 0.0331 |
| GO:0004563 | beta-N-acetylhexosaminidase activity | 2 | 2 | 0.0331 |
| GO:0017136 | NAD-dependent histone deacetylase activity | 2 | 2 | 0.0331 |
| GO:0102148 | N-acetyl-beta-D-galactosaminidase activity | 2 | 2 | 0.0331 |
| GO:0008026 | ATP-dependent helicase activity | 5 | 11 | 0.0344 |
| GO:0005546 | phosphatidylinositol-4,5-bisphosphate binding | 4 | 8 | 0.0409 |
| GO:0004527 | exonuclease activity | 3 | 5 | 0.0449 |
| GO:0030246 | carbohydrate binding | 12 | 40 | 0.0457 |
| **Cellular Component** | | |  |  |
| GO:0042582 | azurophil granule | 4 | 4 | 0.0011 |
| GO:0000790 | nuclear chromatin | 12 | 27 | 0.0014 |
| GO:0031616 | spindle pole centrosome | 3 | 4 | 0.0209 |
| GO:0000786 | nucleosome | 5 | 10 | 0.0223 |
| GO:0042719 | mitochondrial intermembrane space protein transporter complex | 2 | 2 | 0.0333 |
| GO:0001740 | Barr body | 2 | 2 | 0.0333 |
| GO:0072687 | meiotic spindle | 2 | 2 | 0.0333 |
| GO:0034751 | aryl hydrocarbon receptor complex | 2 | 2 | 0.0333 |
| GO:0043196 | varicosity | 2 | 2 | 0.0333 |
| GO:0001931 | uropod | 3 | 5 | 0.0453 |
| **28°C+B3** |  |  |  |  |
| **Biological Process** | |  |  |  |
| GO:0003009 | skeletal muscle contraction | 5 | 7 | 0.0015 |
| GO:0051897 | positive regulation of protein kinase B signaling | 6 | 11 | 0.0034 |
| GO:0010677 | negative regulation of cellular carbohydrate metabolic process | 3 | 3 | 0.0039 |
| GO:1901896 | positive regulation of calcium-transporting ATPase activity | 3 | 3 | 0.0039 |
| GO:0032781 | positive regulation of ATPase activity | 9 | 15 | 0.0055 |
| GO:0050873 | brown fat cell differentiation | 3 | 4 | 0.0138 |
| GO:0006937 | regulation of muscle contraction | 8 | 22 | 0.0147 |
| GO:0071560 | cellular response to transforming growth factor beta stimulus | 8 | 22 | 0.0147 |
| GO:0060048 | cardiac muscle contraction | 6 | 15 | 0.0209 |
| GO:0048193 | Golgi vesicle transport | 12 | 42 | 0.024 |
| GO:1903630 | regulation of aminoacyl-tRNA ligase activity | 2 | 2 | 0.0249 |
| GO:0070836 | caveola assembly | 2 | 2 | 0.0249 |
| GO:1901017 | negative regulation of potassium ion transmembrane transporter activity | 2 | 2 | 0.0249 |
| GO:0009226 | nucleotide-sugar biosynthetic process | 2 | 2 | 0.0249 |
| GO:0071481 | cellular response to X-ray | 2 | 2 | 0.0249 |
| GO:0031571 | mitotic G1 DNA damage checkpoint | 2 | 2 | 0.0249 |
| GO:1904398 | positive regulation of neuromuscular junction development | 2 | 2 | 0.0249 |
| GO:0035414 | negative regulation of catenin import into nucleus | 2 | 2 | 0.0249 |
| GO:0030643 | cellular phosphate ion homeostasis | 2 | 2 | 0.0249 |
| GO:0071378 | cellular response to growth hormone stimulus | 3 | 5 | 0.0305 |
| GO:2000059 | negative regulation of protein ubiquitination involved in ubiquitin-dependent protein catabolic process | 3 | 5 | 0.0305 |
| GO:0035774 | positive regulation of insulin secretion involved in cellular response to glucose stimulus | 3 | 5 | 0.0305 |
| GO:0046627 | negative regulation of insulin receptor signaling pathway | 3 | 5 | 0.0305 |
| GO:0032780 | negative regulation of ATPase activity | 3 | 5 | 0.0305 |
| GO:0034260 | negative regulation of GTPase activity | 3 | 5 | 0.0305 |
| GO:0043392 | negative regulation of DNA binding | 3 | 5 | 0.0305 |
| GO:0008286 | insulin receptor signaling pathway | 7 | 14 | 0.0384 |
| GO:0030518 | intracellular steroid hormone receptor signaling pathway | 6 | 17 | 0.0392 |
| GO:0045913 | positive regulation of carbohydrate metabolic process | 4 | 9 | 0.0399 |
| GO:0000086 | G2/M transition of mitotic cell cycle | 4 | 9 | 0.0399 |
| GO:0097421 | liver regeneration | 5 | 13 | 0.0411 |
| GO:0050772 | positive regulation of axonogenesis | 5 | 13 | 0.0411 |
| GO:0007030 | Golgi organization | 7 | 22 | 0.046 |
| **Molecular Function** | |  |  |  |
| GO:0035259 | glucocorticoid receptor binding | 3 | 4 | 0.013 |
| GO:0001671 | ATPase activator activity | 3 | 4 | 0.013 |
| GO:0008134 | transcription factor binding | 15 | 55 | 0.016 |
| GO:0050431 | transforming growth factor beta binding | 2 | 2 | 0.024 |
| GO:0031730 | CCR5 chemokine receptor binding | 2 | 2 | 0.024 |
| GO:0031014 | troponin T binding | 2 | 2 | 0.024 |
| GO:0005044 | scavenger receptor activity | 3 | 5 | 0.029 |
| GO:0019905 | syntaxin binding | 4 | 9 | 0.037 |
| GO:0016779 | nucleotidyltransferase activity | 4 | 9 | 0.037 |
| GO:0004888 | transmembrane signaling receptor activity | 4 | 9 | 0.037 |
| **Cellular Component** | | |  |  |
| GO:0005887 | integral component of plasma membrane | 12 | 32 | 0.002 |
| GO:0030134 | COPII-coated ER to Golgi transport vesicle | 5 | 9 | 0.0067 |
| GO:0005861 | troponin complex | 2 | 2 | 0.0245 |
| GO:0001741 | XY body | 2 | 2 | 0.0245 |
| GO:0043596 | nuclear replication fork | 3 | 5 | 0.0299 |
| GO:0044295 | axonal growth cone | 3 | 5 | 0.0299 |
| GO:0016459 | myosin complex | 5 | 13 | 0.0399 |

| **Table 3.** Full list of differentially regulated proteins in WAT | | | |
| --- | --- | --- | --- |
| **GeneID** | **Gene name** | **logfc** | **adjpv** |
| **20°C** |  |  |  |
| 117028 | Bin1 | -2.79 | 8.97E-06 |
| 304290 | Kdelr2 | -2.68 | 4.47E-05 |
| 24667 | Ppm1b | -2.65 | 0.000108 |
| 84114 | Agps | -1.35 | 0.000125 |
| 84401 | Puf60 | -2.94 | 0.000753 |
| 300983 | Abhd14b | 0.94 | 0.000791 |
| 29218 | Rcn2 | -2.40 | 0.000854 |
| 290028 | Osgep | -0.94 | 0.002064 |
| 24230 | Tspo | -2.34 | 0.00248 |
| 171452 | Rab3il1 | -2.15 | 0.00629 |
| 84355 | Atox1 | -0.89 | 0.007345 |
| 311428 | RGD1311739 | 1.07 | 0.007461 |
| 25246 | Bsg | 0.84 | 0.007876 |
| 25027 | Slc16a1 | 1.02 | 0.008061 |
| 361999 | Anp32e | -3.03 | 0.00862 |
| 24788 | Sord | 0.81 | 0.01132 |
| 301384 | Hibch | 1.64 | 0.013339 |
| 83576 | Sort1 | 0.96 | 0.014054 |
| 29666 | Psmb6 | 0.72 | 0.017559 |
| 690131 | Hist2h2aa2 | 0.92 | 0.024289 |
| 29428 | Celf2 | -1.64 | 0.024896 |
| 84428 | Dctn4 | 1.81 | 0.031509 |
| 54321 | Cnn3 | -1.91 | 0.032921 |
| 29528 | Vamp3 | 0.68 | 0.034143 |
| 313200 | Hsdl2 | 0.66 | 0.034361 |
| 25287 | Acadl | 0.67 | 0.03642 |
| 83472 | Ugdh | 0.61 | 0.041704 |
| 689284 | Rpl38 | -3.68 | 0.042184 |
| 24383 | Gapdh | -0.72 | 0.046605 |
| 287633 | Lrrc59 | 1.23 | 0.050259 |
| 25604 | Pcmt1 | -0.69 | 0.053192 |
| 681059 | Vps25 | -1.20 | 0.054081 |
| 79223 | Gk | 1.62 | 0.058056 |
| 691947 | Eif3j | -0.97 | 0.062525 |
| 302500 | Mcts1 | 1.98 | 0.065517 |
| 64517 | Thop1 | -1.02 | 0.066596 |
| 84472 | Ilf3 | -3.91 | 0.067026 |
| 65033 | Stx12 | 0.65 | 0.06742 |
| 79131 | Fabp3 | 2.67 | 0.067767 |
| 24172 | Adh1 | 3.28 | 0.075818 |
| 114612 | Ddx39b | -3.63 | 0.076071 |
| 369017 | Krt5 | 2.80 | 0.076106 |
| 362809 | Ptges3 | -0.62 | 0.079947 |
| 54231 | Car2 | 1.69 | 0.081896 |
| 171164 | Gbp2 | -0.65 | 0.08245 |
| 171155 | Hadhb | 0.65 | 0.087263 |
| 25104 | Pc | 0.63 | 0.087972 |
| 294673 | Hexb | 2.37 | 0.088946 |
| 65152 | Pfkm | -2.15 | 0.091004 |
| 24918 | Stat5a | 1.42 | 0.094928 |
| 362115 | Fam129b | 0.99 | 0.106174 |
| 25725 | Prkar1a | -1.02 | 0.106696 |
| 64362 | Des | -0.74 | 0.108631 |
| 29651 | Aldh1a7 | 3.17 | 0.111891 |
| 59108 | Mb | -2.51 | 0.114158 |
| 25698 | Ass1 | 3.41 | 0.115407 |
| 114123 | Sardh | 3.01 | 0.117946 |
| 65151 | Rida | 1.40 | 0.118124 |
| 84357 | Sh3kbp1 | -2.26 | 0.118967 |
| 24248 | Cat | 0.94 | 0.119545 |
| 295284 | Rbm8a | -2.91 | 0.121692 |
| 83527 | Dbnl | -2.21 | 0.122676 |
| 83805 | Src | 1.55 | 0.123643 |
| 363854 | Elavl1 | -0.67 | 0.125734 |
| 29271 | Cfl1 | -1.02 | 0.126378 |
| 307842 | Vac14 | -1.03 | 0.128208 |
| 294568 | Wasf1 | -2.01 | 0.128265 |
| 64526 | Ech1 | 0.62 | 0.128887 |
| 113965 | Hadh | 0.89 | 0.131054 |
| 64679 | Tgm4 | -1.15 | 0.131104 |
| 683313 | LOC683313 | 0.77 | 0.13306 |
| 63864 | Hsd17b10 | 0.86 | 0.138671 |
| 299194 | Ptgr2 | 1.16 | 0.138818 |
| 24307 | Cyp4b1 | -3.32 | 0.143961 |
| 297893 | Hdac1 | -1.83 | 0.151039 |
| 79248 | Abca2 | 1.01 | 0.151314 |
| 300218 | Tuba1c | -3.89 | 0.152971 |
| 81521 | Msn | -0.84 | 0.160485 |
| 24666 | Ppm1a | 1.18 | 0.162655 |
| 102550391 | LOC102550391 | 1.19 | 0.165852 |
| 64533 | Pnpo | -4.31 | 0.166568 |
| 84478 | Ufd1 | 0.87 | 0.172453 |
| 500419 | Rmdn1 | 0.96 | 0.180088 |
| 24360 | Fabp1 | 3.63 | 0.180799 |
| 100125372 | Ces1f | -1.25 | 0.182806 |
| 24284 | Csn1s1 | 2.46 | 0.186806 |
| 299923 | Ndrg1 | 0.63 | 0.187013 |
| 24159 | Acly | 1.25 | 0.19101 |
| 681429 | Rps27l | -0.63 | 0.193334 |
| 84509 | Ran | -0.74 | 0.194937 |
| 246298 | Retsat | 0.97 | 0.195298 |
| 117130 | Grifin | -2.17 | 0.195614 |
| 29474 | Coro1b | -0.63 | 0.196282 |
| 500040 | Tes | 2.33 | 0.198406 |
| 116689 | Ptpn6 | -1.10 | 0.202346 |
| 25499 | Nrdc | 1.35 | 0.202642 |
| 678759 | Ndufa10 | -2.34 | 0.203597 |
| 683788 | Fscn1 | -0.70 | 0.20588 |
| 29459 | Rbbp9 | -1.22 | 0.207521 |
| 25106 | Rgn | 2.76 | 0.208098 |
| 64158 | Tuba1a | 5.02 | 0.218196 |
| 497811 | Xdh | 0.79 | 0.221165 |
| 25371 | Adprh | -2.29 | 0.224889 |
| 306332 | Ap1m1 | 0.71 | 0.227335 |
| 117282 | Hnrnpk | -0.75 | 0.229928 |
| 296710 | Arpc5l | -0.95 | 0.232611 |
| 29443 | Ahcy | -0.90 | 0.23316 |
| 113940 | Gmfg | -1.53 | 0.234475 |
| 24957 | Glul | 1.73 | 0.235712 |
| 60356 | Csad | 1.10 | 0.236915 |
| 24223 | B2m | -0.64 | 0.238063 |
| 363425 | Cav2 | 0.88 | 0.238385 |
| 50671 | Fasn | 1.48 | 0.239956 |
| 64040 | Aldh9a1 | 0.62 | 0.241028 |
| 56781 | Myl1 | -3.64 | 0.241232 |
| 89827 | Ddx39a | -0.78 | 0.244658 |
| **28°C+B3** |  |  |  |
| 83730 | Vamp8 | -3.75 | 6.93E-05 |
| 29521 | Scamp1 | 1.51 | 0.000111 |
| 25116 | Hsd11b1 | 0.92 | 0.000382 |
| 117045 | Eif4e | -0.69 | 0.000942 |
| 25342 | Oxtr | 1.79 | 0.001104 |
| 298566 | C1qa | 0.85 | 0.001124 |
| 445268 | Ufc1 | -0.65 | 0.001133 |
| 78947 | Gcs1 | 0.61 | 0.00204 |
| 266734 | Npas4 | 0.87 | 0.004234 |
| 246303 | Serbp1 | 0.78 | 0.004516 |
| 25139 | Slc2a4 | 1.36 | 0.005653 |
| 619574 | LOC619574 | -1.74 | 0.006446 |
| 64317 | Gpx3 | 1.00 | 0.006558 |
| 84474 | Ddx1 | -0.73 | 0.007131 |
| 24471 | Hspb1 | 1.26 | 0.007327 |
| 170673 | Palm | 2.47 | 0.007749 |
| 313035 | Dnajc8 | -2.09 | 0.008094 |
| 64045 | Glrx | -3.30 | 0.008229 |
| 122799 | Rps25 | -1.04 | 0.009005 |
| 252928 | Timm13 | -0.62 | 0.009087 |
| 25611 | Otc | 0.72 | 0.009208 |
| 54319 | Ezr | -2.37 | 0.009756 |
| 171114 | Ndrg2 | 0.99 | 0.011103 |
| 64306 | Rpl27 | -0.99 | 0.013106 |
| 170520 | Cygb | 0.73 | 0.01364 |
| 85333 | Slc25a4 | 0.74 | 0.014342 |
| 303606 | Ccdc47 | 1.69 | 0.015281 |
| 81520 | Marcksl1 | -2.76 | 0.016098 |
| 116547 | S100a8 | 2.76 | 0.018187 |
| 497009 | Naaa | -3.00 | 0.018457 |
| 494345 | Pdcd10 | -0.80 | 0.020933 |
| 361663 | Lhpp | -1.87 | 0.022049 |
| 25339 | Npr3 | 1.07 | 0.022951 |
| 24233 | C4a | 0.96 | 0.023674 |
| 64152 | Chp1 | 1.02 | 0.025969 |
| 300757 | Hexa | -0.66 | 0.026564 |
| 361051 | Phf11 | -1.84 | 0.026791 |
| 260321 | Fkbp4 | -0.91 | 0.028603 |
| 292148 | Eif3a | -0.71 | 0.030009 |
| 64028 | Tsnax | -3.94 | 0.031875 |
| 100134871 | LOC100134871 | 2.33 | 0.032522 |
| 117259 | Tra2b | -1.82 | 0.032726 |
| 24648 | Serpina1 | 0.98 | 0.033119 |
| 100911615 | LOC100911615 | 1.62 | 0.03415 |
| 292925 | Tsg101 | -1.88 | 0.035767 |
| 300035 | Pycr3 | -0.66 | 0.036528 |
| 360882 | Cadm3 | 1.85 | 0.036758 |
| 308650 | Hnrnph2 | -1.01 | 0.038168 |
| 58927 | Rpl36 | -0.84 | 0.039681 |
| 29635 | Timm44 | -2.58 | 0.041016 |
| 25282 | Cox6a1 | -1.73 | 0.041823 |
| 81504 | Grb2 | -1.13 | 0.043459 |
| 29360 | Selenop | 0.91 | 0.044436 |
| 29286 | Rps17 | -0.67 | 0.046061 |
| 24674 | Ppp3ca | -1.18 | 0.04939 |
| 25524 | Psap | -0.99 | 0.050226 |
| 64352 | Gstm5 | 1.30 | 0.050372 |
| 116698 | Trim28 | -1.28 | 0.050847 |
| 24614 | Orm1 | 1.57 | 0.051012 |
| 171133 | Gcsh | 3.37 | 0.051166 |
| 305679 | Vcl | 0.78 | 0.051276 |
| 25030 | Andpro | 2.99 | 0.051519 |
| 288001 | Kng1 | 0.86 | 0.05194 |
| 64665 | Flot1 | 0.79 | 0.053329 |
| 78958 | Bcam | 1.37 | 0.053481 |
| 681544 | LOC681544 | 0.77 | 0.053624 |
| 24439 | Hagh | 1.13 | 0.056742 |
| 367562 | Gaa | 1.10 | 0.057099 |
| 58827 | Mest | -0.74 | 0.057415 |
| 362401 | Tmem43 | 0.61 | 0.060953 |
| 191574 | Akr1c14 | 0.70 | 0.062102 |
| 294239 | Ddah2 | 0.66 | 0.063782 |
| 24825 | Tf | 0.96 | 0.066528 |
| 65261 | Myo1c | 0.67 | 0.067612 |
| 58917 | Hpx | 0.65 | 0.068548 |
| 85332 | Cavin3 | 0.87 | 0.069924 |
| 690050 | Tpmt | -2.40 | 0.069953 |
| 60581 | Acaca | -3.44 | 0.071 |
| 252929 | Ctsz | -1.05 | 0.072146 |
| 363113 | Syncrip | -0.63 | 0.072433 |
| 296709 | Rpl35 | -0.99 | 0.072466 |
| 300075 | Tomm22 | -0.76 | 0.07444 |
| 100359922 | LOC100359922 | -0.80 | 0.074441 |
| 29236 | Rpsa | -0.68 | 0.074828 |
| 25420 | Cryab | 5.29 | 0.074881 |
| 83712 | Rbbp7 | -0.76 | 0.075646 |
| 25473 | Lamb2 | 0.88 | 0.075929 |
| 29558 | Fcgrt | 0.80 | 0.076078 |
| 56780 | Acpp | -1.89 | 0.078124 |
| 81775 | Rps21 | -1.16 | 0.078693 |
| 29669 | Psma2 | -0.61 | 0.078717 |
| 24440 | Hbb | 2.07 | 0.078843 |
| 360504 | Hba2 | 0.96 | 0.079007 |
| 83783 | Sult1a1 | 0.62 | 0.07972 |
| 117041 | Nln | 3.09 | 0.080074 |
| 25742 | S100b | 1.24 | 0.080926 |
| 300677 | Atp5l | 0.67 | 0.081587 |
| 246233 | Macrod1 | 2.41 | 0.082923 |
| 64507 | Fmod | 2.19 | 0.083367 |
| 116549 | Csnk2a1 | -0.85 | 0.083542 |
| 499782 | Rpl12 | -0.67 | 0.085869 |
| 29283 | Rpl29 | -0.65 | 0.086577 |
| 108348260 | LOC108348260 | -0.73 | 0.092616 |
| 81766 | Rpl18 | -0.67 | 0.096475 |
| 29288 | Rps3a | -0.79 | 0.097091 |
| 500538 | Ybx1 | -0.62 | 0.097152 |
| 360646 | Limd2 | -4.82 | 0.098828 |
| 360471 | Usp7 | -1.03 | 0.103854 |
| 364838 | Reep5 | 1.15 | 0.105331 |
| 57341 | Parva | 0.76 | 0.106336 |
| 171137 | Khsrp | -1.01 | 0.10724 |
| 315218 | Lmf2 | 0.95 | 0.108704 |
| 25035 | Cyb5r3 | 1.02 | 0.110565 |
| 116662 | Ecm1 | 1.41 | 0.111825 |
| 171577 | Epcam | -1.15 | 0.112584 |
| 307779 | Rbmxrtl | -0.96 | 0.115081 |
| 307947 | Set | -2.08 | 0.117698 |
| 60571 | Mybbp1a | -1.40 | 0.11861 |
| 64031 | Pdcd4 | -1.13 | 0.119102 |
| 65984 | Aacs | -3.23 | 0.120908 |
| 362855 | Rtcb | -0.61 | 0.121081 |
| 56611 | Anxa2 | 0.97 | 0.122294 |
| 296596 | Rpl7a | -0.70 | 0.12345 |
| 114113 | Pafah1b3 | -1.02 | 0.12414 |
| 369016 | Myadm | 0.73 | 0.124691 |
| 28298 | Rpl32 | -1.06 | 0.125753 |
| 450225 | Krt10 | 1.17 | 0.127176 |
| 59114 | Slc9a3r1 | -1.09 | 0.12891 |
| 25757 | Cpt1a | 3.07 | 0.12931 |
| 317381 | Ccdc22 | 5.59 | 0.129886 |
| 85255 | Hacl1 | -2.44 | 0.130084 |
| 302562 | Plp2 | 0.71 | 0.131025 |
| 362631 | Rpl11 | -0.75 | 0.132254 |
| 691531 | Rps28 | -0.87 | 0.133683 |
| 79224 | Serpind1 | 0.66 | 0.134675 |
| 192276 | Coro7 | -1.16 | 0.136285 |
| 81681 | Lss | 1.38 | 0.141036 |
| 55939 | Apom | 0.63 | 0.141553 |
| 81729 | Rpl10a | -0.98 | 0.142368 |
| 360626 | Krt19 | -2.95 | 0.142817 |
| 291434 | Rpl17 | -0.66 | 0.143666 |
| 305343 | Pds5a | -3.37 | 0.145504 |
| 25419 | Crp | 0.87 | 0.147213 |
| 290651 | Isyna1 | -1.36 | 0.148286 |
| 360854 | Arpc5 | -0.76 | 0.151956 |
| 80846 | Hnrnpl | -1.05 | 0.153574 |
| 300079 | Rpl3 | -0.81 | 0.153618 |
| 50664 | Gnao1 | 0.91 | 0.153753 |
| 29491 | Itsn1 | -3.57 | 0.153827 |
| 113936 | Cpb2 | 0.93 | 0.154271 |
| 100360522 | LOC100360522 | -1.22 | 0.155548 |
| 79256 | Hnrnpd | -1.69 | 0.15639 |
| 360576 | Tusc5 | 1.02 | 0.158706 |
| 308384 | Sae1 | -1.99 | 0.159931 |
| 296654 | Gsn | 0.87 | 0.160988 |
| 361673 | Ifitm3 | -1.59 | 0.165339 |
| 29389 | Tnni2 | -4.01 | 0.165619 |
| 29671 | Psma4 | -0.73 | 0.166824 |
| 65204 | Cnn1 | 2.94 | 0.168593 |
| 301252 | Hsp90ab1 | -0.75 | 0.170109 |
| 24786 | Sod1 | -0.92 | 0.170236 |
| 81008 | Itga7 | 1.13 | 0.170372 |
| 117557 | Tpm3 | -0.84 | 0.173177 |
| 497794 | Mug1 | 0.61 | 0.174553 |
| 58952 | Cpq | 0.62 | 0.175249 |
| 114499 | Hdgf | -0.82 | 0.178124 |
| 65137 | Ruvbl1 | -1.53 | 0.178135 |
| 25010 | Scgb2a1 | 4.76 | 0.179267 |
| 117042 | Rpl6 | -0.79 | 0.180849 |
| 287191 | Rars | -0.82 | 0.181074 |
| 25686 | Gnai1 | 0.63 | 0.183971 |
| 294853 | Krt18 | -2.59 | 0.185246 |
| 286938 | Gimap4 | -1.75 | 0.186195 |
| 29648 | Nudc | -0.86 | 0.186791 |
| 85496 | Enpp1 | -1.97 | 0.18985 |
| 83502 | Cdh1 | -1.92 | 0.19445 |
| 25330 | Lipe | 1.13 | 0.195171 |
| 83510 | Lypla2 | -0.91 | 0.195635 |
| 29473 | Aoc3 | 0.70 | 0.195941 |
| 117280 | Hnrnpu | -1.07 | 0.196818 |
| 116655 | Hnrnpm | -1.12 | 0.200116 |
| 25126 | Stat5b | -0.87 | 0.200585 |
| 24366 | Fgb | 0.86 | 0.200904 |
| 108350501 | LOC108350501 | -1.35 | 0.202062 |
| 64205 | Rplp0 | -0.79 | 0.205181 |
| 79116 | Apex1 | -3.25 | 0.206758 |
| 25368 | Adk | -0.72 | 0.210856 |
| 290641 | Rpl18a | -0.77 | 0.211728 |
| 170724 | Anp32b | -1.02 | 0.212806 |
| 297699 | Strap | -0.70 | 0.213497 |
| 100362830 | LOC100362830 | -0.77 | 0.218146 |
| 24968 | Psmb8 | -0.84 | 0.219647 |
| 252922 | Pzp | 0.96 | 0.223427 |
| 291983 | Psmb10 | -1.07 | 0.22378 |
| 116685 | Lmnb1 | -1.10 | 0.224812 |
| 361512 | Ehd2 | 1.19 | 0.227089 |
| 25269 | Pvalb | 2.17 | 0.227186 |
| 108348062 | LOC108348062 | -2.51 | 0.228587 |
| 29563 | Crabp2 | -3.47 | 0.238336 |
| 64347 | Sncg | 1.49 | 0.23862 |
| 25292 | Apoc1 | 2.25 | 0.239531 |
| 309187 | Atl3 | 0.84 | 0.240149 |
| 290644 | Ifi30 | -0.70 | 0.240249 |
| 24346 | Ces1c | 0.91 | 0.243417 |
| 103690821 | LOC103690821 | -0.66 | 0.245662 |
| 293692 | Ehd1 | 1.02 | 0.247676 |

| **Table 4. Full list of GO terms enriched in WAT** | | | | |
| --- | --- | --- | --- | --- |
| **goId** | **goName** | **countDE** | **countAll** | **pv_elim** |
| **20°C** | | | | |
| **Biological Process** | | | | |
| GO:0030330 | DNA damage response, signal transduction by p53 class mediator | 5 | 6 | 0.0022 |
| GO:0048711 | positive regulation of astrocyte differentiation | 3 | 3 | 0.0023 |
| GO:0071498 | cellular response to fluid shear stress | 3 | 3 | 0.0023 |
| GO:0032780 | negative regulation of ATPase activity | 3 | 3 | 0.0023 |
| GO:0051607 | defense response to virus | 5 | 9 | 0.003 |
| GO:0001822 | kidney development | 11 | 35 | 0.0036 |
| GO:0050731 | positive regulation of peptidyl-tyrosine phosphorylation | 7 | 17 | 0.0037 |
| GO:0002244 | hematopoietic progenitor cell differentiation | 3 | 4 | 0.0081 |
| GO:0045577 | regulation of B cell differentiation | 3 | 4 | 0.0081 |
| GO:0042130 | negative regulation of T cell proliferation | 3 | 4 | 0.0081 |
| GO:0000077 | DNA damage checkpoint | 3 | 4 | 0.0081 |
| GO:0002763 | positive regulation of myeloid leukocyte differentiation | 3 | 4 | 0.0081 |
| GO:0071803 | positive regulation of podosome assembly | 3 | 4 | 0.0081 |
| GO:0016477 | cell migration | 26 | 127 | 0.0104 |
| GO:0034314 | Arp2/3 complex-mediated actin nucleation | 8 | 13 | 0.0124 |
| GO:0010592 | positive regulation of lamellipodium assembly | 4 | 8 | 0.0133 |
| GO:0120033 | negative regulation of plasma membrane bounded cell projection assembly | 4 | 8 | 0.0133 |
| GO:0051289 | protein homotetramerization | 8 | 26 | 0.0147 |
| GO:1902743 | regulation of lamellipodium organization | 8 | 13 | 0.0164 |
| GO:2000279 | negative regulation of DNA biosynthetic process | 4 | 5 | 0.017 |
| GO:0000245 | spliceosomal complex assembly | 4 | 5 | 0.017 |
| GO:2000573 | positive regulation of DNA biosynthetic process | 6 | 17 | 0.017 |
| GO:1902570 | protein localization to nucleolus | 2 | 2 | 0.0173 |
| GO:0032480 | negative regulation of type I interferon production | 2 | 2 | 0.0173 |
| GO:0071362 | cellular response to ether | 2 | 2 | 0.0173 |
| GO:0034616 | response to laminar fluid shear stress | 2 | 2 | 0.0173 |
| GO:0006499 | N-terminal protein myristoylation | 2 | 2 | 0.0173 |
| GO:0097484 | dendrite extension | 2 | 2 | 0.0173 |
| GO:0035855 | megakaryocyte development | 2 | 2 | 0.0173 |
| GO:0099601 | regulation of neurotransmitter receptor activity | 2 | 2 | 0.0173 |
| GO:0035970 | peptidyl-threonine dephosphorylation | 2 | 2 | 0.0173 |
| GO:2000394 | positive regulation of lamellipodium morphogenesis | 2 | 2 | 0.0173 |
| GO:0010870 | positive regulation of receptor biosynthetic process | 2 | 2 | 0.0173 |
| GO:0031954 | positive regulation of protein autophosphorylation | 2 | 2 | 0.0173 |
| GO:0060740 | prostate gland epithelium morphogenesis | 2 | 2 | 0.0173 |
| GO:2000601 | positive regulation of Arp2/3 complex-mediated actin nucleation | 2 | 2 | 0.0173 |
| GO:1902463 | protein localization to cell leading edge | 2 | 2 | 0.0173 |
| GO:2000107 | negative regulation of leukocyte apoptotic process | 3 | 5 | 0.0184 |
| GO:0048011 | neurotrophin TRK receptor signaling pathway | 3 | 5 | 0.0184 |
| GO:0042475 | odontogenesis of dentin-containing tooth | 3 | 5 | 0.0184 |
| GO:0051701 | interaction with host | 8 | 27 | 0.0187 |
| GO:0030203 | glycosaminoglycan metabolic process | 4 | 9 | 0.0215 |
| GO:0050792 | regulation of viral process | 8 | 28 | 0.0232 |
| GO:0051091 | positive regulation of DNA binding transcription factor activity | 7 | 23 | 0.0236 |
| **Molecular Function** | |  |  |  |
| GO:0051287 | NAD binding | 11 | 31 | 0.0013 |
| GO:0008144 | drug binding | 9 | 29 | 0.0101 |
| GO:0005001 | transmembrane receptor protein tyrosine phosphatase activity | 2 | 2 | 0.0178 |
| GO:0005521 | lamin binding | 3 | 5 | 0.0191 |
| GO:0042393 | histone binding | 5 | 13 | 0.021 |
| GO:0033613 | activating transcription factor binding | 3 | 6 | 0.0345 |
| GO:0003857 | 3-hydroxyacyl-CoA dehydrogenase activity | 3 | 6 | 0.0345 |
| GO:0045296 | cadherin binding | 22 | 113 | 0.0365 |
| GO:0004028 | 3-chloroallyl aldehyde dehydrogenase activity | 2 | 3 | 0.0486 |
| GO:0071933 | Arp2/3 complex binding | 2 | 3 | 0.0486 |
| GO:0004854 | xanthine dehydrogenase activity | 2 | 3 | 0.0486 |
| GO:0003785 | actin monomer binding | 2 | 3 | 0.0486 |
| GO:0046912 | transferase activity, transferring acyl groups, acyl groups converted into alkyl on transfer | 2 | 3 | 0.0486 |
| GO:0005324 | long-chain fatty acid transporter activity | 2 | 3 | 0.0486 |
| GO:0005540 | hyaluronic acid binding | 2 | 3 | 0.0486 |
| **Cellular Component** | | |  |  |
| GO:0005884 | actin filament | 9 | 24 | 0.0021 |
| GO:0032993 | protein-DNA complex | 5 | 11 | 0.0088 |
| GO:0002102 | podosome | 7 | 14 | 0.013 |
| GO:0016607 | nuclear speck | 8 | 26 | 0.0146 |
| GO:0031209 | SCAR complex | 2 | 2 | 0.0172 |
| GO:0042611 | MHC protein complex | 2 | 2 | 0.0172 |
| GO:0005687 | U4 snRNP | 2 | 2 | 0.0172 |
| GO:0005856 | cytoskeleton | 49 | 238 | 0.0284 |
| GO:0030054 | cell junction | 41 | 216 | 0.0291 |
| GO:0005681 | spliceosomal complex | 9 | 22 | 0.0311 |
| GO:0031258 | lamellipodium membrane | 3 | 6 | 0.033 |
| GO:0071437 | invadopodium | 3 | 6 | 0.033 |
| GO:0005912 | adherens junction | 29 | 162 | 0.0408 |
| GO:0000407 | pre-autophagosomal structure | 2 | 3 | 0.0471 |
| GO:0002080 | acrosomal membrane | 2 | 3 | 0.0471 |
| **28°C+B3** |  |  |  |  |
| **Biological Process** | |  |  |  |
| GO:0006953 | acute-phase response | 7 | 12 | 0.0035 |
| GO:0000381 | regulation of alternative mRNA splicing, via spliceosome | 7 | 12 | 0.0035 |
| GO:0034113 | heterotypic cell-cell adhesion | 8 | 12 | 0.0061 |
| GO:0070528 | protein kinase C signaling | 4 | 5 | 0.0063 |
| GO:0015671 | oxygen transport | 4 | 5 | 0.0063 |
| GO:1901741 | positive regulation of myoblast fusion | 3 | 3 | 0.0077 |
| GO:0070934 | CRD-mediated mRNA stabilization | 3 | 3 | 0.0077 |
| GO:0007566 | embryo implantation | 6 | 12 | 0.0179 |
| GO:0040007 | growth | 29 | 103 | 0.0209 |
| GO:0071345 | cellular response to cytokine stimulus | 25 | 86 | 0.0211 |
| GO:0009059 | macromolecule biosynthetic process | 84 | 332 | 0.0239 |
| GO:0070293 | renal absorption | 3 | 4 | 0.0261 |
| GO:0071392 | cellular response to estradiol stimulus | 3 | 4 | 0.0261 |
| GO:0042993 | positive regulation of transcription factor import into nucleus | 3 | 4 | 0.0261 |
| GO:0048821 | erythrocyte development | 3 | 4 | 0.0261 |
| GO:0046597 | negative regulation of viral entry into host cell | 3 | 4 | 0.0261 |
| GO:0044319 | wound healing, spreading of cells | 3 | 4 | 0.0261 |
| GO:0043516 | regulation of DNA damage response, signal transduction by p53 class mediator | 3 | 4 | 0.0261 |
| GO:0031953 | negative regulation of protein autophosphorylation | 3 | 4 | 0.0261 |
| GO:2000648 | positive regulation of stem cell proliferation | 3 | 4 | 0.0261 |
| GO:1900087 | positive regulation of G1/S transition of mitotic cell cycle | 3 | 4 | 0.0261 |
| GO:0042273 | ribosomal large subunit biogenesis | 8 | 15 | 0.027 |
| GO:0030032 | lamellipodium assembly | 6 | 13 | 0.0278 |
| GO:0006281 | DNA repair | 10 | 22 | 0.0281 |
| GO:0032103 | positive regulation of response to external stimulus | 11 | 31 | 0.0287 |
| GO:0051241 | negative regulation of multicellular organismal process | 33 | 123 | 0.0288 |
| GO:0042255 | ribosome assembly | 8 | 20 | 0.0291 |
| GO:0042307 | positive regulation of protein import into nucleus | 7 | 11 | 0.0307 |
| GO:0070670 | response to interleukin-4 | 5 | 10 | 0.0308 |
| GO:0072659 | protein localization to plasma membrane | 15 | 47 | 0.0309 |
| GO:0071407 | cellular response to organic cyclic compound | 33 | 103 | 0.0311 |
| GO:0034114 | regulation of heterotypic cell-cell adhesion | 4 | 7 | 0.0317 |
| GO:0042755 | eating behavior | 4 | 7 | 0.0317 |
| GO:2001235 | positive regulation of apoptotic signaling pathway | 9 | 24 | 0.0325 |
| GO:0019915 | lipid storage | 7 | 17 | 0.0346 |
| GO:0051090 | regulation of DNA binding transcription factor activity | 10 | 28 | 0.0348 |
| GO:0090316 | positive regulation of intracellular protein transport | 15 | 32 | 0.0354 |
| GO:0045861 | negative regulation of proteolysis | 21 | 67 | 0.0377 |
| GO:0060088 | auditory receptor cell stereocilium organization | 2 | 2 | 0.039 |
| GO:0032415 | regulation of sodium:proton antiporter activity | 2 | 2 | 0.039 |
| GO:0061158 | 3'-UTR-mediated mRNA destabilization | 2 | 2 | 0.039 |
| GO:2001014 | regulation of skeletal muscle cell differentiation | 2 | 2 | 0.039 |
| GO:0050884 | neuromuscular process controlling posture | 2 | 2 | 0.039 |
| GO:2001026 | regulation of endothelial cell chemotaxis | 2 | 2 | 0.039 |
| GO:0019731 | antibacterial humoral response | 2 | 2 | 0.039 |
| GO:0071386 | cellular response to corticosterone stimulus | 2 | 2 | 0.039 |
| GO:0006388 | tRNA splicing, via endonucleolytic cleavage and ligation | 2 | 2 | 0.039 |
| GO:2000047 | regulation of cell-cell adhesion mediated by cadherin | 2 | 2 | 0.039 |
| GO:0006407 | rRNA export from nucleus | 2 | 2 | 0.039 |
| GO:2001137 | positive regulation of endocytic recycling | 2 | 2 | 0.039 |
| GO:0019886 | antigen processing and presentation of exogenous peptide antigen via MHC class II | 2 | 2 | 0.039 |
| GO:0060355 | positive regulation of cell adhesion molecule production | 2 | 2 | 0.039 |
| GO:0002523 | leukocyte migration involved in inflammatory response | 2 | 2 | 0.039 |
| GO:0031643 | positive regulation of myelination | 2 | 2 | 0.039 |
| GO:0072697 | protein localization to cell cortex | 2 | 2 | 0.039 |
| GO:0010642 | negative regulation of platelet-derived growth factor receptor signaling pathway | 2 | 2 | 0.039 |
| GO:0080111 | DNA demethylation | 2 | 2 | 0.039 |
| GO:0052405 | negative regulation by host of symbiont molecular function | 2 | 2 | 0.039 |
| GO:1905063 | regulation of vascular smooth muscle cell differentiation | 2 | 2 | 0.039 |
| GO:0030643 | cellular phosphate ion homeostasis | 2 | 2 | 0.039 |
| GO:0010757 | negative regulation of plasminogen activation | 2 | 2 | 0.039 |
| GO:0071763 | nuclear membrane organization | 2 | 2 | 0.039 |
| GO:0060669 | embryonic placenta morphogenesis | 2 | 2 | 0.039 |
| GO:0035176 | social behavior | 2 | 2 | 0.039 |
| GO:0018065 | protein-cofactor linkage | 2 | 2 | 0.039 |
| GO:0015886 | heme transport | 2 | 2 | 0.039 |
| GO:0050427 | 3'-phosphoadenosine 5'-phosphosulfate metabolic process | 2 | 2 | 0.039 |
| GO:0002925 | positive regulation of humoral immune response mediated by circulating immunoglobulin | 2 | 2 | 0.039 |
| GO:0010998 | regulation of translational initiation by eIF2 alpha phosphorylation | 2 | 2 | 0.039 |
| GO:0007156 | homophilic cell adhesion via plasma membrane adhesion molecules | 2 | 2 | 0.039 |
| GO:0000387 | spliceosomal snRNP assembly | 2 | 2 | 0.039 |
| GO:0007194 | negative regulation of adenylate cyclase activity | 2 | 2 | 0.039 |
| GO:1904401 | cellular response to Thyroid stimulating hormone | 2 | 2 | 0.039 |
| GO:0005984 | disaccharide metabolic process | 2 | 2 | 0.039 |
| GO:0000447 | endonucleolytic cleavage in ITS1 to separate SSU-rRNA from 5.8S rRNA and LSU-rRNA from tricistronic rRNA transcript (SSU-rRNA, 5.8S rRNA, LSU-rRNA) | 2 | 2 | 0.039 |
| GO:0000461 | endonucleolytic cleavage to generate mature 3'-end of SSU-rRNA from (SSU-rRNA, 5.8S rRNA, LSU-rRNA) | 2 | 2 | 0.039 |
| GO:0000463 | maturation of LSU-rRNA from tricistronic rRNA transcript (SSU-rRNA, 5.8S rRNA, LSU-rRNA) | 2 | 2 | 0.039 |
| GO:0038089 | positive regulation of cell migration by vascular endothelial growth factor signaling pathway | 2 | 2 | 0.039 |
| GO:1903533 | regulation of protein targeting | 6 | 14 | 0.0408 |
| GO:0000165 | MAPK cascade | 22 | 78 | 0.0414 |
| GO:0022409 | positive regulation of cell-cell adhesion | 9 | 25 | 0.0423 |
| GO:0006396 | RNA processing | 31 | 75 | 0.0432 |
| GO:0045596 | negative regulation of cell differentiation | 19 | 66 | 0.0463 |
| GO:0032092 | positive regulation of protein binding | 7 | 18 | 0.0474 |
| GO:0051053 | negative regulation of DNA metabolic process | 5 | 11 | 0.0475 |
| GO:0007188 | adenylate cyclase-modulating G-protein coupled receptor signaling pathway | 5 | 11 | 0.0475 |
| GO:0034381 | plasma lipoprotein particle clearance | 5 | 11 | 0.0475 |
| GO:0008154 | actin polymerization or depolymerization | 17 | 42 | 0.0483 |
| **Molecular Function** | |  |  |  |
| GO:0003735 | structural constituent of ribosome | 24 | 62 | 0.00037 |
| GO:0005344 | oxygen carrier activity | 4 | 5 | 0.00657 |
| GO:0003730 | mRNA 3'-UTR binding | 8 | 18 | 0.01548 |
| GO:0003682 | chromatin binding | 10 | 25 | 0.01623 |
| GO:0140097 | catalytic activity, acting on DNA | 5 | 9 | 0.01908 |
| GO:0042162 | telomeric DNA binding | 3 | 4 | 0.02688 |
| GO:0045294 | alpha-catenin binding | 3 | 4 | 0.02688 |
| GO:0004527 | exonuclease activity | 3 | 4 | 0.02688 |
| GO:0003723 | RNA binding | 73 | 281 | 0.02734 |
| GO:0019825 | oxygen binding | 4 | 7 | 0.03278 |
| GO:0031720 | haptoglobin binding | 2 | 2 | 0.03976 |
| GO:0001091 | RNA polymerase II basal transcription factor binding | 2 | 2 | 0.03976 |
| GO:0001105 | RNA polymerase II transcription coactivator activity | 2 | 2 | 0.03976 |
| GO:0015926 | glucosidase activity | 2 | 2 | 0.03976 |
| GO:0005055 | laminin receptor activity | 2 | 2 | 0.03976 |
| GO:0005089 | Rho guanyl-nucleotide exchange factor activity | 2 | 2 | 0.03976 |
| GO:0045159 | myosin II binding | 2 | 2 | 0.03976 |
| GO:0043395 | heparan sulfate proteoglycan binding | 2 | 2 | 0.03976 |
| GO:0046790 | virion binding | 2 | 2 | 0.03976 |
| GO:0004888 | transmembrane signaling receptor activity | 7 | 13 | 0.04822 |
| GO:0070851 | growth factor receptor binding | 5 | 11 | 0.04938 |
| **Cellular Component** | | |  |  |
| GO:0022625 | cytosolic large ribosomal subunit | 15 | 30 | 0.00018 |
| GO:0016323 | basolateral plasma membrane | 14 | 32 | 0.00164 |
| GO:0005833 | hemoglobin complex | 3 | 3 | 0.00782 |
| GO:0005903 | brush border | 10 | 23 | 0.00812 |
| GO:0030864 | cortical actin cytoskeleton | 9 | 20 | 0.00918 |
| GO:0016327 | apicolateral plasma membrane | 3 | 4 | 0.02666 |
| GO:0044451 | nucleoplasm part | 18 | 58 | 0.0269 |
| GO:0005637 | nuclear inner membrane | 5 | 10 | 0.03167 |
| GO:0035770 | ribonucleoprotein granule | 11 | 32 | 0.03804 |
| GO:0097225 | sperm midpiece | 2 | 2 | 0.03953 |
| GO:0032426 | stereocilium tip | 2 | 2 | 0.03953 |
| GO:0061827 | sperm head | 2 | 2 | 0.03953 |
| GO:0070937 | CRD-mediated mRNA stability complex | 2 | 2 | 0.03953 |
| GO:0072669 | tRNA-splicing ligase complex | 2 | 2 | 0.03953 |
| GO:0071204 | histone pre-mRNA 3'end processing complex | 2 | 2 | 0.03953 |
| GO:0034451 | centriolar satellite | 2 | 2 | 0.03953 |
| GO:0030686 | 90S preribosome | 2 | 2 | 0.03953 |
